# Human ORFeome expression in *S. cerevisiae* to better understand extracellular vesicle biology

**DOI:** 10.64898/2026.09.05.749614

**Authors:** Joseph Trani, Sara Daneshi, Aashiq Hussain Kachroo, Christopher Leonard Brett

## Abstract

Extracellular vesicles (EVs) mediate intercellular communication by all organisms studied, from bacteria to yeast to man. Yet evolutionarily conserved mechanisms governing aspects of fundamental EV biology remain enigmatic. To address this, we sought to establish *Saccharomyces cerevisiae* (baker’s yeast) as a model by identifying ectopically expressed human proteins sorted into yeast EVs. Using an optimized pooled cloning method, we inserted >13,000 human open reading frames (ORFs) upstream of an EGFP tag within yeast expression plasmids. After transformation into *S. cerevisiae*, we confirmed expression of 3,288 EGFP-tagged human proteins with diverse cellular expression levels and subcellular localizations. Heat stress triggered release of intact, lipid-bound, EGFP-positive small EVs from all transformant pools. Proteomic analysis identified 292 human proteins within EV samples, including canonical human EV biomarkers. Over 70% had yeast orthologs also found in yeast EVs suggesting conserved sorting mechanisms. Protein–protein interaction network analysis linked these EV cargoes to ESCRT-associated pathways. Finally, validation of seven candidates showed that DEF3A, ANXA2 and CLIC1 were enriched in yeast EVs. This study establishes an omics-compatible synthetic biology framework to humanize yeast EVs, begins to uncover conserved cargo sorting mechanisms, and supports future engineering of designer EVs.

**GRAPHICAL ABSTRACT and TOC BLURB:** 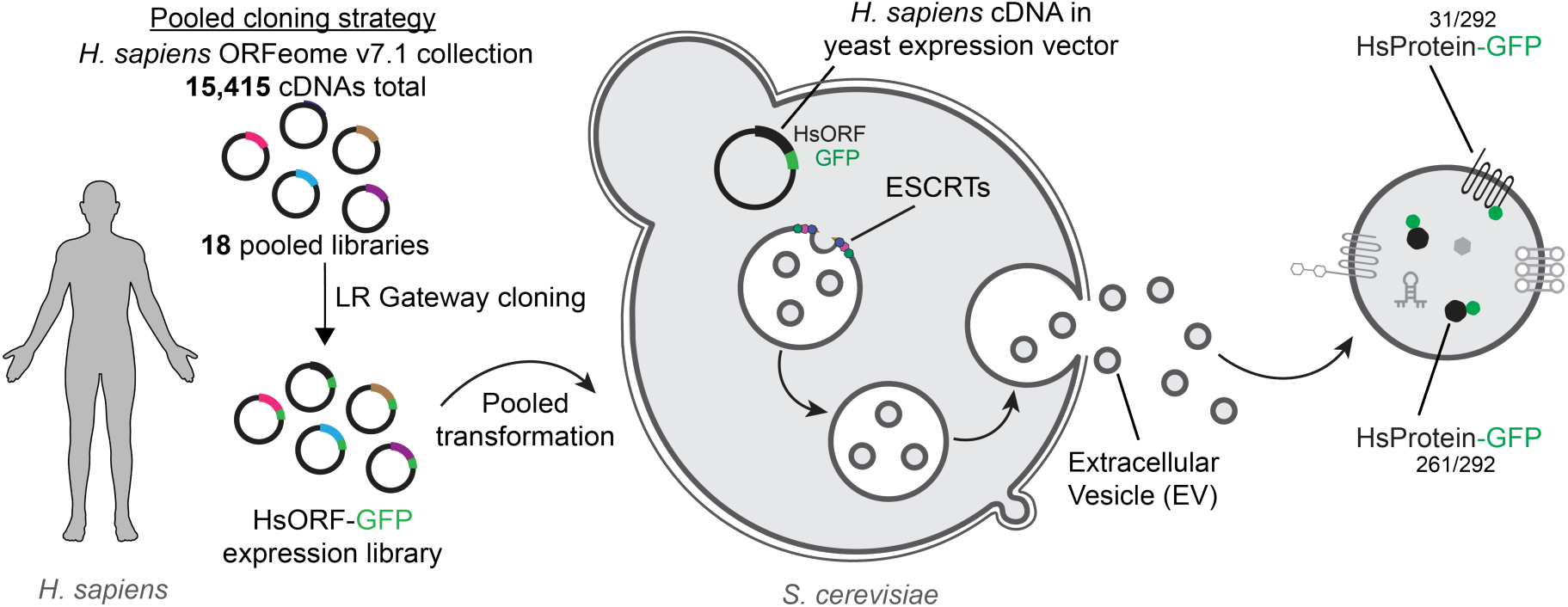

Over 13,000 human open reading frames were expressed in *Saccharomyces cerevisiae* to study fundamental extracellular vesicle (EV) biology. Results suggest evolutionary conservation of EV cargo protein sorting pathways and identify human protein scaffolds to engineer yeast EVs for broad applications.

## INTRODUCTION

Extracellular vesicles (EVs) are membrane-bound nanocarriers that transport proteins, nucleic acids, and lipids between cells as part of a conserved intercellular communication system (Mathieu et al., 2019; van Niel et al., 2018). EVs are biosynthesized by donor cells and secretion is regulated by physiological cues and environmental stressors, including heat shock, oxidative stress, and hypoxia (Bister et al., 2020; Gebremedhn et al., 2020; Hedlund et al., 2011). Once released, EVs are selectively recognized by and deliver their bioactive cargo to recipient cells via fusion with the plasma membrane fusion or endosome membranes after endocytosis. Cargo biomolecules then drive signaling thought to mediate diverse physiology, e.g. modulating immune responses and influencing tissue development (Bahram Sangani et al., 2021; Kalluri & LeBleu, 2020; Ragni, 2025). In pathological contexts, EV-based signaling is implicated in tumour progression, remodeling and metastasis, and in prion protein propagation underlying neurodegenerative diseases (Asai et al., 2015; Hu et al., 2016; Lucotti et al., 2022). Despite advances in knowledge of EV (patho)physiology, the molecular mechanisms that mediate most aspects of EV biology – biogenesis, cargo loading, secretion, recognition, fusion – remain unresolved.

Beyond their roles in physiology and disease, EVs are attractive drug delivery vehicles because they are biologically derived, membrane-bound nanoparticles capable of transporting diverse molecular cargo between cells. These properties could, in principle, be engineered to protect therapeutic molecules, alter biodistribution, promote selective delivery, or introduce new biological functions. However, these same capabilities depend on poorly defined processes, including cargo selection, vesicle biogenesis, secretion, target-cell recognition, and intracellular delivery. This mechanistic uncertainty limits the rational engineering of EV-based therapeutics and remains a major barrier to realizing their full potential as predictable drug delivery systems.

Although most research focuses on human health, EV-based communication is an evolutionarily ancient process observed across phyla, i.e. bacteria, protozoa, yeasts, plants, animals (Brown et al., 2015; Gill et al., 2019; Rizzo et al., 2020). The budding yeast *Saccharomyces cerevisiae*, a genetically tractable eukaryote, has long been a cornerstone for dissecting mechanisms responsible for fundamental eukaryotic processes such as vesicle trafficking, proteostasis, transcription and cell cycle regulation (Feyder et al., 2015; Sampaio-Marques et al., 2018; Vanderwaeren et al., 2022). *S. cerevisiae* was also instrumental in the discovery and characterization of the endosomal sorting complexes required for transport (ESCRTs) and ALIX, which are thought to mediate small EV or exosome biogenesis in humans (Baietti et al., 2012; Hurley & Hanson, 2010; Pashkova et al., 2013). While EV release from *S. cerevisiae* has been documented for over a decade (Mencher et al., 2020; Oliveira, et al., 2010; Rodrigues & Nimrichter, 2022), early work largely catalogued vesicle biomolecular content without revealing the underlying mechanism(s) responsible for EV biology. More recently, we demonstrated that EVs shared by *S. cerevisiae* contribute to thermotolerance (Logan et al., 2024), setting the stage for establishing *S. cerevisiae* as a model to study the basis of fundamental EV biology in detail.

Furthermore, ectopically expressing human genes in *S. cerevisiae* (or “humanizing” them) is a powerful approach used to study evolutionary cell biology and to better understand effects of disease alleles on cellular processes, e.g. mutant α-synuclein, mutant huntingtin, proteasome components (Kachroo et al., 2015, 2022; Laval et al., 2023). Amenable to complex genetic engineering, *S. cerevisiae* is often used as a chassis organism in the field of synthetic biology that merges principles in engineering and molecular biology to create novel biological systems (Rainha et al., 2020). With this in mind, we recently completed a proof-of-concept study that optimized synthetic biology framework and modular cloning system called EVclo to genetically engineer *S. cerevisiae* EVs (Bouffard et al., 2026). We used EVclo to demonstrate that a small peptide derived from human EV proteins (called ExoSignal; Ferreira et al., 2022) can be sorted into yeast EVs. Finally, *S. cerevisiae* is used to manufacturer biological drugs, offering an opportunity to scale up bioproduction of future EV-based technologies (Nielsen, 2013).

Here, we sought to further develop *S. cerevisiae* as a platform to study EV biology and engineer EVs for broad applications by leveraging its experimental advantages to uncover evolutionarily conserved mechanisms responsible for EV protein sorting. We also aimed to determine whether individual human proteins could be selectively incorporated into yeast EVs, providing a foundation to later explore their functions in isolation and identify candidate scaffolds for EV engineering. We hypothesized that if *S. cerevisiae* and humans share EV cargo-sorting mechanisms, then a fraction of known human EV proteins should be selectively enriched in yeast EVs when ectopically expressed. To test this hypothesis, we designed an unbiased screening experiment that unites our EVclo system (Bouffard et al., 2026) with an optimized pooled cloning method to introduce up to 15,415 different human open reading frames (ORFs) tagged with EGFP into *S. cerevisiae*. Human proteins within EVs collected from pooled transformants were then identified by mass spectrometry and subjected to bioinformatic analysis. We reasoned that this approach would reveal conserved cargo-selection principles, demonstrate selective EV incorporation, and identify synthetic scaffolds - including EV-enriched human proteins without yeast orthologs - for future engineering strategies.

## RESULTS

### A pooled cloning method to express the human ORFeome in *S. cerevisiae*

To initiate our study, we first developed and optimized a pooled cloning method to express the human ORFeome library version 7.1 (Center for Cancer Systems Biology hORFeome v7.1) in *S. cerevisiae*. This collection contains 18,414 human ORFs (HsORFs), representing 15,415 unique genes, encoded within Gateway compatible donor vectors (Lamesch et al., 2007). To maximize representation of HsORF expression, we divided this large collection into 18 pools that each contained between 420 to 940 cDNAs. In theory, this approach should improve confidence of downstream HsORF detection, by pooled next-generation DNA sequencing or protein identification by mass spectrometry, and help reduce HsORF expression bias later. Each pool of HsORFs, flanked by AttL sties within donor plasmids, was then subjected to Gateway LR recombination to incorporate them into a destination vector, containing AttR sites, from our modular EVclo system (Bouffard et al., 2026). Resulting expression plasmids each encode a transcriptional unit (TU) – containing a strong constitutive *S. cerevisiae* promoter (PTDH3) followed by a single human ORF (HsORF) flanked by AttB sites, an in-frame EGFP tag for fusion protein detection, and an efficient terminator (T_ADH1_) – as well as a high copy replication system (2µ) and a URA3 marker for plasmid maintenance and yeast transformant selection (Fig. 1a).

**Figure 1.**
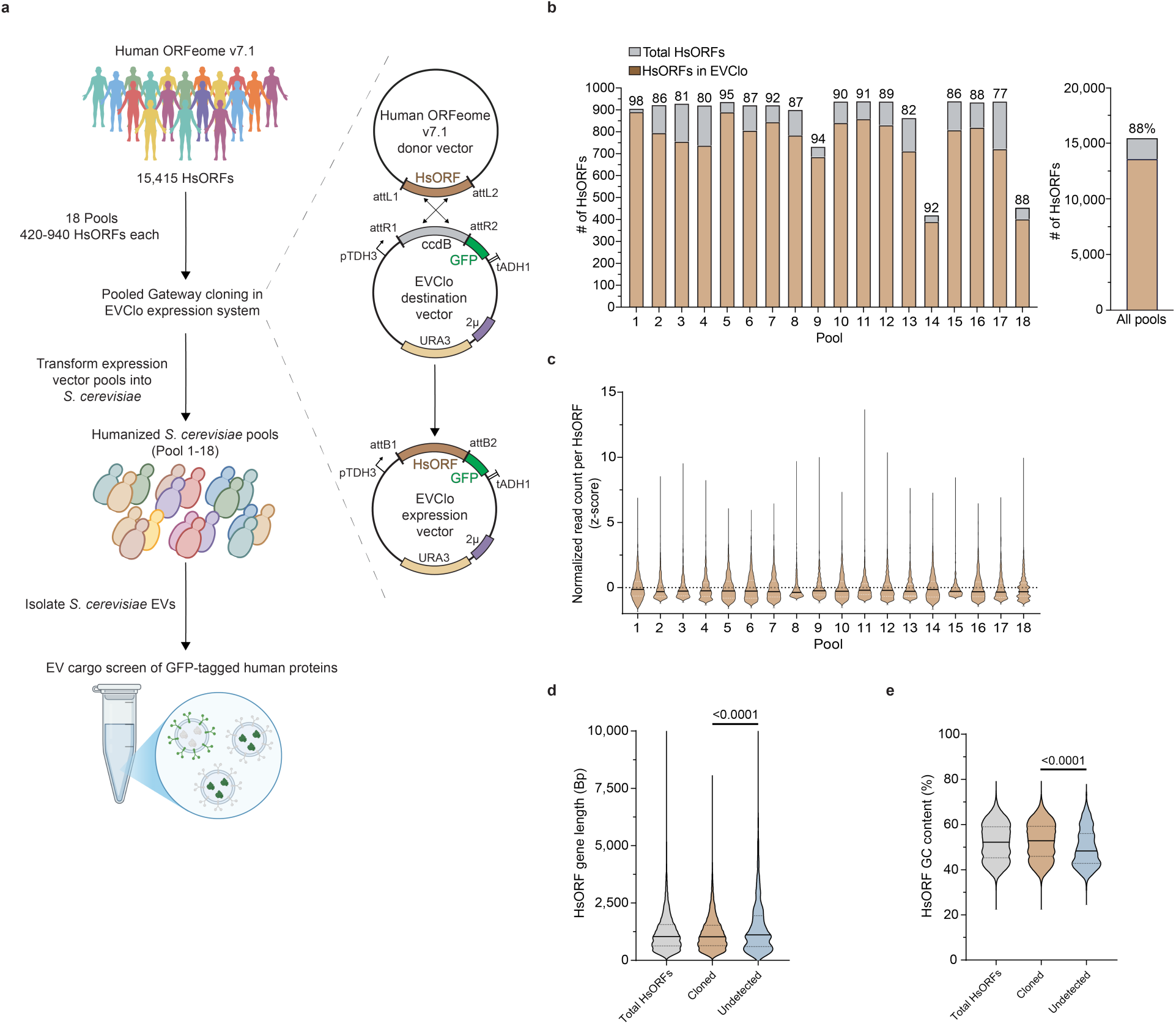
Pooled Gateway cloning enables broad representation of the human ORFeome in *Saccharomyces cerevisiae*. **a,** Schematic of the pooled cloning and screening workflow. The human ORFeome v7.1 HsORF collection was divided into 18 pools and hORFS were transferred by Gateway LR recombination from donor vectors into the EVClo destination vector. The resulting EVClo expression plasmids encode a constitutive *S. cerevisiae* TDH3 promoter (pTDH3), a single human ORF (HsORF), an in-frame green fluorescent protein (GFP) tag, an *S. cerevisiae* ADH1 terminator (tADH1), a 2µ high-copy origin of replication, and a URA3 selectable marker. Expression pools were transformed into *S. cerevisiae*, EVs were isolated, and GFP-tagged human EV cargo proteins were identified. **b,** Graph depicting pooled cloning efficiency of hORFs into EVClo expression vectors. Grey bars indicate total input HsORFs per pool, and brown bars indicate HsORFs detected after pooled cloning. Values above bars denote the percentage of input HsORFs detected in each pool. Across all 18 pools, 13,614 of 15,415 total HsORFs were recovered (88%). **c,** Violin plots showing the distribution of z-score–normalized read counts per HsORF for detected HsORFs within each pool, calculated from raw read counts and normalized within each pool. Center lines indicate the median; dotted lines indicate quartiles. **d,** Violin plots comparing HsORF length distributions for the total input library, cloned HsORFs, and undetected HsORFs. Cloned HsORFs were defined as HsORFs detected in Nanopore-sequenced EVClo expression pools; undetected HsORFs were present in the input library but not detected after pooled cloning. Center lines indicate the median; dotted lines indicate quartiles. Cloned versus undetected HsORFs were compared using a two-tailed Mann–Whitney test. **e,** Violin plots comparing GC content distributions for the total input library, cloned HsORFs, and uncloned HsORFs. Center lines indicate the median; dotted lines indicate quartiles. Cloned versus undetected HsORFs were compared using a two-tailed Mann–Whitney test.

To quantify cloning efficiency and assess HsORF coverage, we performed Oxford Nanopore long-read sequencing of expression plasmid DNA from each *E. coli*–amplified pool prior to yeast transformation. Pool-specific success rates ranged from 77% to 98%, but across all 18 pools, we inserted 13,614 unique HsORFs into expression plasmids, representing 88% of the original hORFeome v. 7.1 library (Fig. 1b; gene IDs shown in Supplementary Information 1). To evaluate representation bias within each pool, we calculated Z-score–normalized read distributions for all detected HsORFs. Most pools of HsORF containing expression plasmids exhibited unimodal distributions centered around zero, indicating minimal skewing in HsORF abundance and uniform representation across the pools (Fig. 1c; DNA fragment abundance shown in Supplementary Information 1). However, we noticed that HsORF gene length affected cloning efficiency, as cloned HsORFs were significantly shorter than undetected HsORFs (Fig. 1d). Also, HsORF GC content was significantly higher in cloned HsORFs compared to undetected HsORFs (Fig. 1e). These findings suggest that this pooled cloning method is somewhat efficient – although it favours shorter, GC-rich sequences – and retains library complexity while minimizing stochastic dropout or overamplification of specific hORFs, all critical for unbiased downstream screening studies.

### hORFeome expression produces diverse human proteins in *S. cerevisiae*

We next transformed wild type *S. cerevisiae* with these 18 pools of HsORF-containing expression plasmids. Ectopic human gene expression in yeast does not always lead to protein translation for many reasons (most theoretical and unresolved). Thus, to help maximize the number of cells expressing human proteins, transformants were selected based on uracil auxotrophy as well as detectable GFP fluorescence by FACS (fluorescence-activated cell sorting) to enrich cells carrying the plasmid and expressing GFP-tagged human proteins, respectively. To confirm cellular expression of GFP-tagged human proteins, we examined pooled cultures by flow cytometry (Fig. 2a) and found that they contained between 39% and 72% (or 54% ± 10% across pools) detectable GFP-positive cells (Fig. 2b). Cellular GFP fluorescence intensity was variable across pools and cells (Fig. 2a and c). This was expected given that GFP-positive cells contained different HsORFs that may be expressed at different levels due to (i) sequence variability, despite being driven by a common promotor, and (ii) different subcellular distributions.

**Figure 2.**
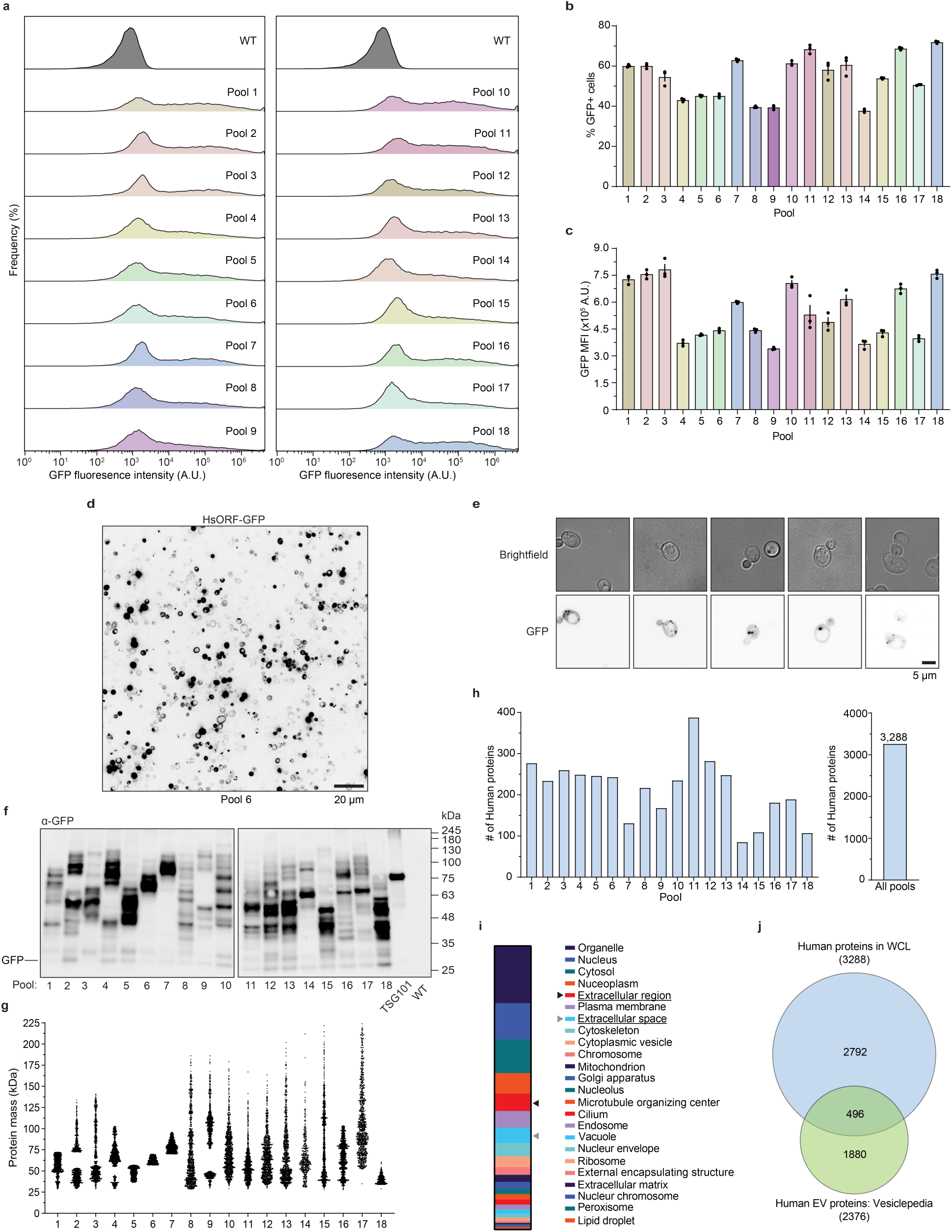
(previous page). Expression of GFP-tagged human proteins in pooled *S. cerevisiae*. **a,** Flow cytometry histograms showing GFP fluorescence distributions for WT cells and each of the 18 pooled *S. cerevisiae* cultures expressing GFP-tagged human ORFs. **b,** Percentage of GFP-positive cells detected in each pool by flow cytometry. **c,** Mean Fluorescence Intensity (MFI) of GFP-positive cells in each pool measured by flow cytometry. **d,** Representative inverted GFP fluorescence micrograph of cells from pool 6. Scale bar, 20 μm. **e,** Representative brightfield and GFP fluorescence micrographs of individual cells from multiple pools. Scale bar, 5 μm. **f,** Anti-GFP immunoblot of whole cell lysates (WCLs) prepared from each pooled yeast culture. WT cells served as a negative control, and cells expressing TSG101–GFP served as a positive control; the predicted molecular mass of TSG101–GFP is ∼75 kDa. **g,** Predicted molecular mass distribution of all human proteins in each pool tagged with GFP (26.9 kDa). Each point represents one human-GFP fusion protein. **h,** Number of human proteins identified by mass spectrometry in each pool and across all pools. A total of 3,288 unique human proteins were identified in WCLs. **i,** Gene Ontology cellular compartment annotation of the 3,288 human proteins identified in WCLs. Compartments that potentially included EVs are highlighted with arrow heads (Extracellular region, Extracellular space). **j,** Venn diagram comparing human proteins identified in WCLs with proteins annotated in Vesiclepedia under the selected inclusion criteria. Data in **b,c** are mean ± S.E.M. from n = 3 biological replicates.

The latter was confirmed by imaging pooled cultures using confocal microscopy (Fig. 2d and Supplementary Fig. 1), whereby GFP was observed in diverse cellular compartments which was expected when expressing the human proteome in yeast cells. As free GFP localizes exclusively to the cytoplasm (Carminati & Stearns, 1998), observed patterns suggest that the sequence/structural features required for compartmentalization of human proteins (fused to GFP) may also be recognized in *S. cerevisiae* despite extreme evolutionary divergence. Although presumptive, GFP signal is observed in distinct puncta near vacuole membranes indicative of MVBs, presumed sites of ESCRT-mediated EV biogenesis (Fig. 2e; Wollert & Hurley, 2010), suggesting that perhaps some GFP-tagged human proteins are sorted into yeast extracellular vesicles.

The diversity of cellular distributions observed also suggests the translation of intact GFP-tagged human protein fusions. To confirm, we conducted Western blot analysis of whole cell lysates prepared from the 18 pooled yeast cultures subjected to GFP immunoprecipitation (Fig. 2f). We found that all pools exhibited distinct patterns of GFP-reactive bands across a range of molecular weights, which were larger than GFP alone (26.8 kDa); samples prepared from yeast expressing GFP-tagged human TSG101 (70.8 kDa), an EV biomarker (Théry, et al., 2018; Uniprot Q99816), served as a positive control, while the wildtype parent strain served as a negative control. We next calculated predicted sizes of human proteins tagged to GFP found in each pool (Fig. 3g) and found that these patterns were similar to band patterns observed by immunoblotting, further confirming widespread translation of the hORFeome in yeast.

**Figure 3.**
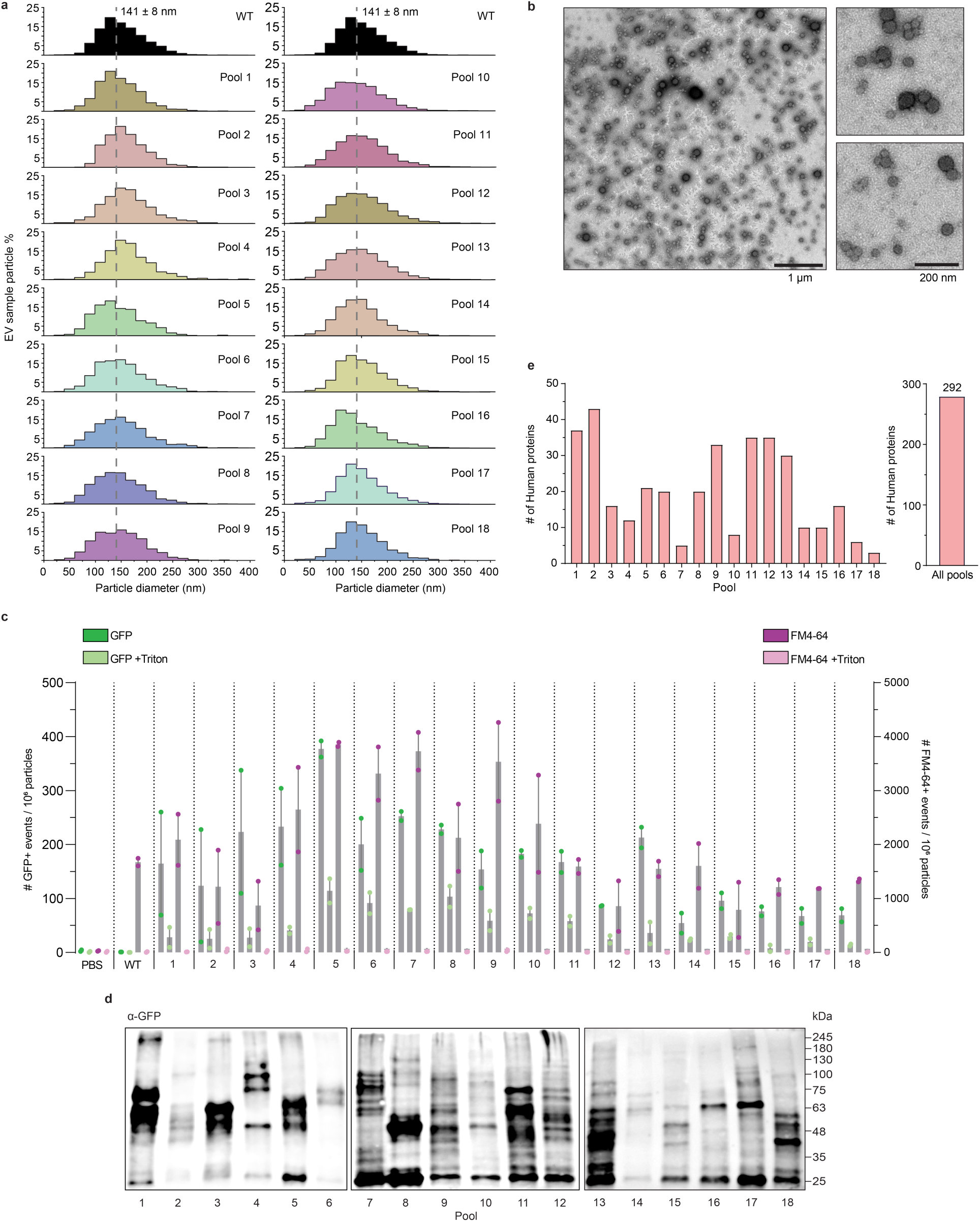
(previous page). Detection of GFP-tagged human proteins in EV samples collected from pooled *S. cerevisiae* transformants. **a,** Representative particle size distributions of EV samples isolated from WT cells or each of the 18 pooled *S. cerevisiae* cultures expressing GFP-tagged human ORFs measured using NTA. The dashed line indicates the mean particle diameter of WT (141 ± 8 nm). **b,** Representative transmission electron micrographs of EV samples isolated from Pool 13. Scale bars, 1 μm and 200 nm. **c,** Nano-flow cytometry analysis of EV samples from each pool, normalized using NTA particle counts. Number of GFP-positive particles (green) and FM4-64-positive particles (magenta) are shown. Lighter shaded data points indicate samples treated with Triton X-100. PBS and WT served as a negative controls. Bars indicate means ± S.E.M.. n = 2 biological replicates. **d,** Anti-GFP immunoblot of EV samples collected from each pool of transformants. **e,** Number of human proteins identified by mass spectrometry in GFP-immunopurified fractions from EV samples isolated from each pool and across all pools. A total of 279 unique human proteins were identified.

To identify human proteins and quantify expression breadth, we performed mass spectrometry-based proteomic profiling on GFP-immunopurified yeast lysates from each pool. We identified between 85 to 388 proteins in each pool of transformants representing 3,288 total unique human proteins across all pools (Fig. 2h; see Supplementary Information 2). This subset of the human proteome includes proteins found in extracellular compartments (that includes EVs) and all intracellular compartments (Fig. 2i), in support of our findings from micrographic analysis (Fig. 2d and e). Importantly, this subset includes 496 human proteins previously found in human EV samples (Fig. 2j), representing 63 of the top 100 listed on Vesiclepedia (a trusted online repository of EV proteomic datasets; (Pathan, Fonseka, et al., 2019)). Thus, although 24% of HsORFs in EVclo were translated in *S. cerevisiae*, many human EV proteins are represented in this subpopulation validating further study of these pooled humanized yeast strains to help uncover conserved mechanisms of EV protein sorting.

### Some human proteins are selectively incorporated into yeast EVs

To identify human proteins sorted into *S. cerevisiae* EVs, we subjected cultures of pooled transformants to heat stress, an established trigger of EV release from yeast and human cells (Logan et al., 2024). We then collected EVs from the extracellular medium by ultrafiltration and size exclusion chromatography and first confirmed their presence by nanoparticle tracking analysis (NTA; Fig. 3a). We found that each pool of transformants secreted 8 to 18 particles per cell with unimodal size distributions centered between 135 and 172 nm in diameter (Supplementary Fig. 2). The average particle diameter of all pools, 141 ± 8, is consistent with the size of yeast and human small EVs, or exosomes, reported by others (Oliveira, et al., 2010; Yáñez-Mó et al., 2015). The average number of purified particles recovered was 12 ± 3 particles per cell, comparable to previous reports using similar yeast EV isolation workflows (Bouffard et al., 2026; Logan et al., 2024). While particle yields varied between pools, they remained broadly consistent with expected yeast EV production, suggesting that pooled HsORF expression did not grossly perturb small EV biosynthesis or release under these conditions.

To confirm that these samples contained EVs, we examined them using transmission electron microscopy (TEM; Fig. 3b) and observed round structures resembling vesicles with diameters of ∼ 100 nm consistent with data from NTA (Fig. 3a) and published micrographs of small EVs (Bouffard et al., 2026; Colombo et al., 2014; Théry et al., 2006). To demonstrate that observed particles were lipid-bound, we stained EVs in samples with the lipophilic dye FM4-64 and conducted nano-flow cytometry (Fig. 3e; Corso et al., 2019; Coumans et al., 2017; Welsh et al., 2023). We confirmed the presence of FM4-64 positive membrane-bound vesicles. We further demonstrated this by treating samples with the detergent Triton-X100 to dissolve vesicles and found complete collapse of FM4-64 signals across all pools (Fig. 3e). To show that EVs contained GFP-tagged human proteins, we also examined particle GFP-fluorescence by nano-flow cytometry (Fig. 3e). We found that a fraction of particles were GFP+ and this signal collapsed with Triton-X100 treatment. Together, these data demonstrate that all pools contained detergent-sensitive, lipid-bound vesicles and detectable GFP-positive particle populations, consistent with the incorporation of GFP-tagged human proteins into small EVs released by *S. cerevisiae*.

Next, we conformed that GFP-human protein fusions are present in yeast EV samples by conducting Western blot analysis (Fig. 3d and Supplementary Fig. 2). We observed distinct patterns of GFP-reactive bands above the size of GFP alone (26.8 kDa) in all 18 pools, representing a subset of proteins observed in whole cell lysates (see Fig. 2f). With confidence that some GFP-tagged human proteins are within yeast EVs, we subjected EV samples from all 18 pools to GFP-immunoaffinity purification and conducted proteomics analysis by mass spectrometry on bound fractions. After peptide spectra were searched against a combined *S. cerevisiae* and human protein databases, we identified 292 unique human proteins in small EV samples collected from all pools (listed in Supplementary Information 2). Notably, nearly all proteins identified were also reported in human EVs (212 of 292) including 31 of the top 100 human proteins found in diverse samples (listed on Vesiclepedia; Pathan, Fonseka, et al., 2019; Pathan, Keerthikumar, et al., 2019), e.g. the human small EV biomarker and scaffold ANAX2 (Supplementary Fig. 2). Thus, because only a fraction (∼9%) of all human proteins expressed in yeast cells were present in their EVs, we conclude that some human proteins can be selectively sorted into small EVs by *S. cerevisiae*.

### Small EV protein sorting mechanisms may be evolutionarily conserved

With this new data in hand, we next conducted bioinformatic analysis to better understand the properties of human proteins found in yeast EV samples and to possibly help uncover a conserved sorting mechanism. We first assessed orthology by determining if genes encoding the 292 human proteins observed in yeast EVs have orthologs encoded in the *S. cerevisiae* genome (Fig. 4a; Supplementary Information 3). We find that 202 human proteins (69%) have orthologs, which is significantly higher than the human ORFeome (52%). Moreover, of these human proteins with *S. cerevisiae* orthologs, we find that 90 of 202 (45%) of their yeast counterparts are also identified within EVs (Logan et al., 2024) and these conserved EV protein cargoes represent 20 of the top 100 most abundant and consistently identified EV proteins (Figure 4a; Supplementary Information 3). This subset of 20 proteins is shown in Table 2 and likely represent small EV cargoes sorted by conserved mechanisms shared between yeast and man underlying fundamental EV biology. Consistent with this interpretation, human proteins recovered from yeast EVs were significantly enriched for Vesiclepedia Top 100 proteins compared to those detected in whole-cell lysates, with 31 of 292 EV proteins overlapping this list compared to 63 of 3,288 whole-cell proteins (10.62% versus 1.92%; Fig. 4b and c). Thus, human proteins incorporated into yeast EVs are disproportionately represented among canonical human EV cargoes, further supporting the idea that *S. cerevisiae* selectively packages human proteins through conserved EV-sorting mechanisms.

**Figure 4.**
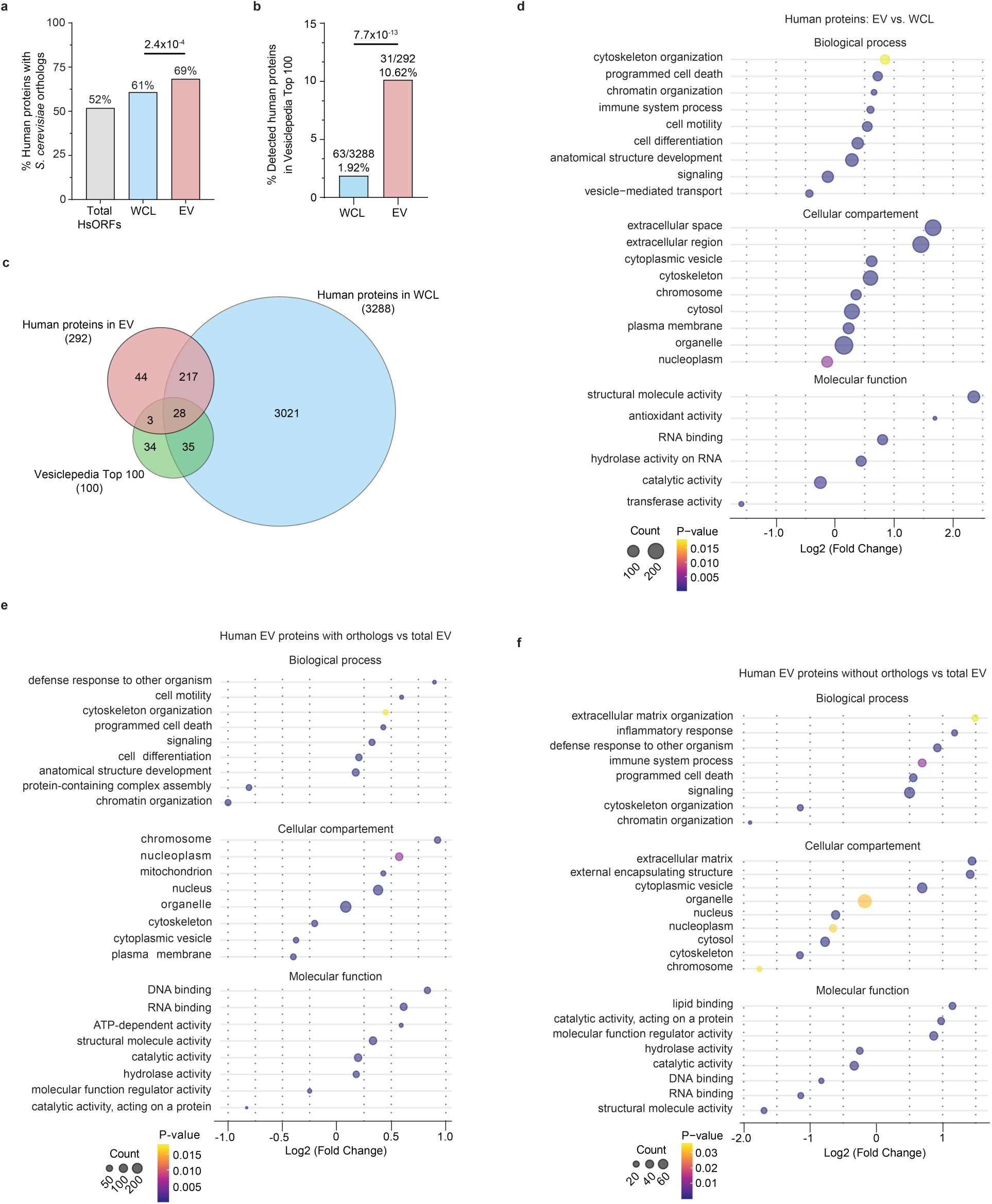
(previous page). Bioinformatic analysis of human proteins identified in *S. cerevisiae* EV samples. **a,** Percentage of human proteins with *S. cerevisiae* orthologs for total HsORFs screened, human proteins identified in yeast WCLs, and human proteins identified in EV samples. Ortholog relationships were assigned using the DRSC Integrative Orthology Prediction Tool (DIOPT). Statistical comparison shown for WCL versus EV is by Fisher’s exact test. **b,** Percentage of human proteins identified in WCLs and EV samples represented in the Vesiclepedia Top 100 protein set. Statistical comparison shown for WCL versus EV is by Fisher’s exact test. **c,** Venn diagram showing the overlap between human proteins identified in EV samples and in WCLs, and the Vesiclepedia Top 100 set. **d,** Gene Ontology (GO) enrichment and depletion analysis of human proteins identified in EVs relative to human proteins identified in WCLs, shown for Biological Process, Cellular Compartment, and Molecular Function categories. **e,** GO enrichment and depletion analysis of EV sample human proteins with *S. cerevisiae* orthologs relative to all human proteins identified in EV samples, shown for Biological Process, Cellular Compartment, and Molecular Function categories. **f,** GO enrichment and depletion analysis of EV sample human proteins without *S. cerevisiae* orthologs relative to all human proteins identified in EV samples, shown for Biological Process, Cellular Compartment, and Molecular Function categories. GO term annotations were obtained using g:Profiler. Dot size indicates protein count and color indicates *P*-value. Only categories with significant enrichment or depletion are shown.

To gain insight into potential functions of all 292 human proteins found in yeast EVs, we performed Gene Ontology (GO) analysis and compared results to analysis of the 3,288 human proteins identified in yeast whole-cell lysates and all 15,415 proteins encoded in the hORFeome v7.1 library (Supplementary Fig. 3). We examined three GO terms: Biological Process, Molecular Function, and Cellular Compartment and found that EV protein distributions were different in each category compared to cellular proteins and the hORFeome, suggesting EV protein selection as anticipated. Yeast EVs were significantly enriched for human proteins associated with the extracellular space in support of our hypothesis, as well as cytoskeleton, nucleoplasm, and chromatin a pattern mirroring features of human EV proteomes (Fig. 4d) (Kowal et al., 2016; Lim et al., 2021; Valadi et al., 2007). Molecular functions such as structural molecule activity, RNA binding, hydrolase activity, and antioxidant activity were overrepresented in the yeast EV human proteome. Biological processes such as cytoskeleton organization, signaling, chromatin organization, immune response, and programmed cell death were also overrepresented consistent with reported roles of human EVs in intercellular communication, immune cell signaling and apoptosis (Lim et al., 2021; Yáñez-Mó et al., 2015). Altogether, this analysis supports the idea that small EV protein sorting mechanism(s) are evolutionarily conserved.

To better understand fundamental EV biology, we repeated this GO analysis on human proteins with *S. cerevisiae* orthologs to possibly reveal evolutionarily ancient EV functions (Fig. 4e). Enriched biological processes include cytoskeleton organization, defense response to other organisms, and cell motility. Overrepresented molecular functions include RNA and DNA binding, ATP-dependent enzymatic activity, and structural molecule function. Together, these results suggest roles in altering cellular morphology, gene regulation and proteostasis, consistent with reports of EV function shared across phyla (Yáñez-Mó et al., 2015). Also, we sought to assess functionalities gained by yeast EVs through humanization by analyzing human proteins that lacked *S. cerevisiae* orthologs. As shown in Figure 4f, enriched biological processes include extracellular matrix organization, inflammatory responses, and immune system processes. Although these functions are not native to yeast, their recovery in yeast EVs suggests that human-specific cargoes may be accommodated by conserved sorting pathways when expressed in *S. cerevisiae* (Parreira et al., 2021; Russell et al., 2020). Moreover, these results demonstrate that *S. cerevisiae* may be humanized to gain synthetic functionalities, not found in nature, possibly supporting future therapeutic applications.

To possibly elucidate a mechanism of cargo sorting, we first examined polypeptide amino acid sequences to potentially identify a common sorting sequence or domain but were unable to find any valid candidates (data not shown). Instead, we considered presumed pathways for EV biogenesis that are shared between mammals and yeasts. We therefore focused on small EVs because reported *S. cerevisiae* EVs are typically within the submicron range, and release of larger vesicle classes may be limited by yeast morphology and cell wall architecture. Haploid *S. cerevisiae* cells are approximately 4–5 μm in diameter, similar in scale to apoptotic bodies, which are typically 1–5 μm, while microvesicles are commonly defined as plasma membrane-derived vesicles of ∼0.1–1 μm. Moreover, estimated *S. cerevisiae* cell wall pore diameters of ∼200–400 nm suggest that the wall likely restricts passage of larger vesicles, consistent with evidence that cell wall integrity influences yeast EV release (Kakarla et al., 2020; Oliveira et al., 2010; Sherman, 2002; Ståhl et al., 2019; Zhao et al., 2019). Thus, we focused on the ESCRT (endosomal sorting complexes required for transport) machinery because orthologs are found in humans and *S. cerevisiae* (where they were first discovered) and it is thought to mediate small EV biogenesis in mammals (Colombo et al., 2014; Hurley, 2015).

To determine potential interactions between all human proteins within EV samples identified in our study and ESCRTs, we constructed a protein-protein interaction map based on STRING databases and STRING-based confidence scoring (Figure 5; see Supplementary Information 4). We also examined ESCRT accessory proteins ALIX, a human EV biomarker, and subunits of the Vps4-Vta complex but found no interactions. Otherwise, 199 of 292 human EV proteins are reported to interact with at least one other protein in this group. Importantly, we observed 6 proteins that directly interact with ESCRT subunits implicated in protein cargo recognition (Henne et al., 2011). We speculate that these proteins may mediate protein selection by ESCRTs and they seem to represent three entry pathways:

**Figure 5.**
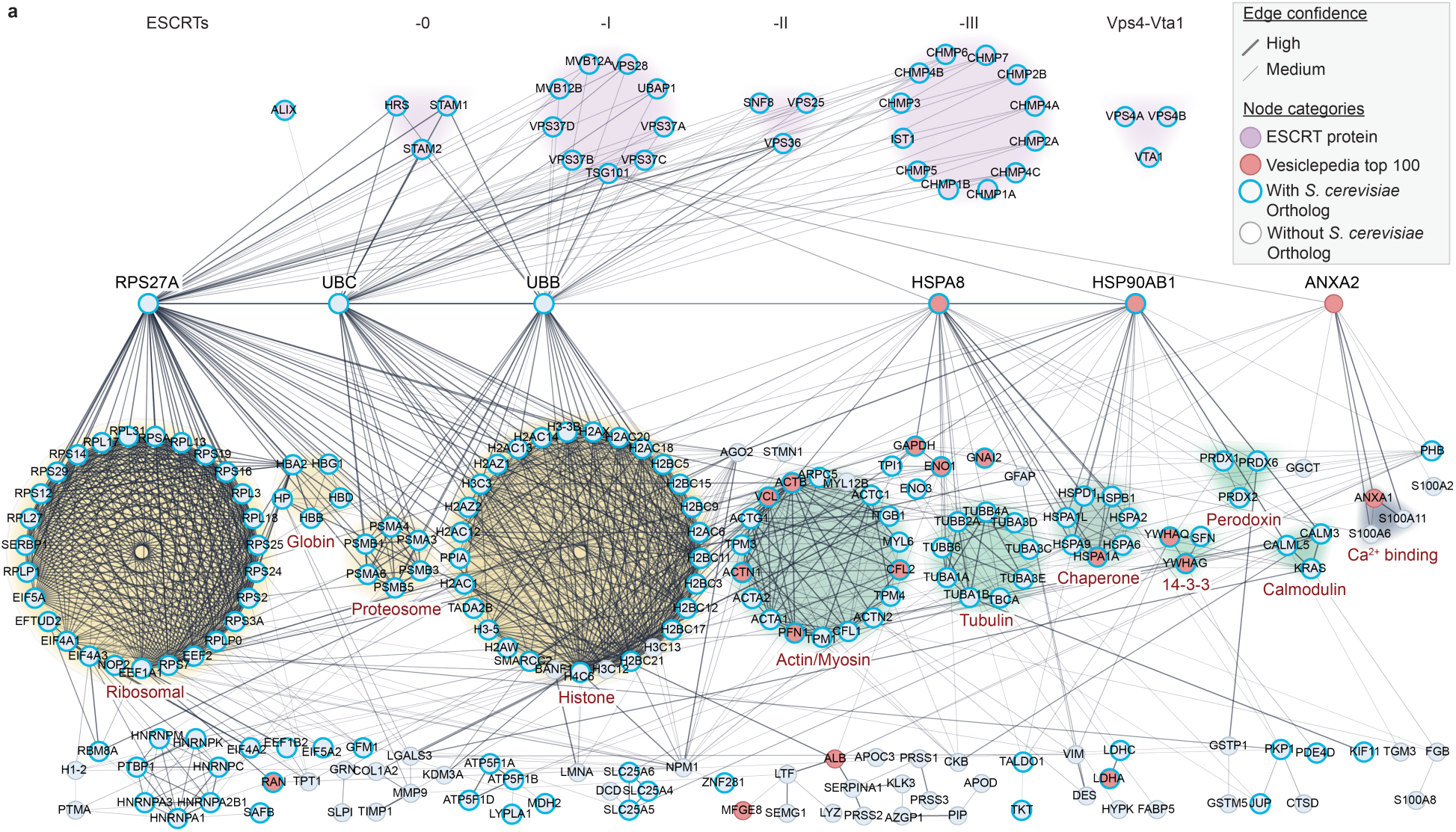
Protein–protein interaction network linking human proteins found in yeast EV samples to the ESCRT machinery. Protein–protein interaction network generated using the STRING extension in Cytoscape software, showing 199 EV sample human proteins linked directly or indirectly to ESCRT-associated proteins or each other. Proteins are represented by nodes indicating ESCRT subunits for each complex (ESCRT-0, -1, -2 and -3, and Vps4-Vta1; magenta), Vesiclepedia Top 100 human protein (red), human proteins with (blue) or without (grey) *S. cerevisiae* orthologs. High and medium confidence protein interactions are represented by edges. Protein clusters demonstrating high connectivity are shown and represent ribosome components, globins, proteosome subunits, histones, actin/myosin, tubulin, chaperones, 14-3-3 proteins, perodoxins, calmodulin, and Ca^2+^ binding proteins associated with Annexin 2 (ANXA2). These clusters are coloured based on high connectivity with ubiquitin orthologs (RPS27A, UBC, UBB; yellow), heat shock proteins (HSPA8, HSP90AB1; green) or annexin A2 (ANXA2, yellow) which directly interact with ESCRT subunits. Human ESCRT proteins were not identified in our screen.

First, many ESCRT subunits interact with ubiquitin paralogs UBC, RPS27A and UBB. Ubiquitins label cargoes for recognition by ESCRTs, e.g. Hrs in ESCRT-0 and TSG101 in ESCRT-I, and thus are critical for protein sorting (Henne et al., 2011). Here we find that they interact with clustered components of the ribosome (27 interacting proteins), globulin (5), proteasome (6), and histones (30) also found within EVs. Although speculative, this suggests that proteins in these clusters may be loaded into EVs through direct ubiquitylation or ubiquitylation of a binding partner within the complex. Nearly all human proteins (96 %) in this subnetwork have yeast orthologs suggesting that this presumed sorting mechanism may be conserved. Given that ubiquitylation is a major signal for proteasomal degradation and protein quality control, we speculate that this subnetwork may represent a disposal route linked to cellular proteostasis, particularly because ribosomal and proteasomal proteins are also implicated (Amm et al., 2014; Glickman & Ciechanover, 2002; Sung et al., 2016).

Second, the ESCRT-I subunit TSG101 implicated in cargo selection interacts with heat shock protein chaperones HSPA8 and HSP90AB1. These form complexes with other heat shock chaperones (7 interacting proteins) and directly interact with clustered components of the cytoskeleton including actin and myosin (18) and tubulin (9), 14-3-3 scaffolds (3), perodoxin (3) and calmodulin-binding proteins (3). In addition, they interact with AGO2, which is thought to mediate RNA loading into EVs (McKenzie et al., 2016), and a cluster that includes GAPDH, ENO1 and TPI1 involved in glycolysis but have other moonlighting functions (Shegay et al., 2023; Sriram et al., 2005). Again, nearly all (99%) proteins in this subnetwork have *S. cerevisiae* orthologs and notably, many of them (14 of 43) are found on the top 100 EV protein list (Vesiclepedia; Pathan, Fonseka, et al., 2019), including HSPA8 and HSP90AB1 themselves, making this a strong candidate as a potential conserved sorting mechanism. Given that heat shock protein chaperones are involved with protein refolding or disassembling protein aggregates, and both HSP8A and HSP90AB1 seem to be ubiquitylated, it is possible that this pathway mediates protein degradation. However, functions of proteins in this network include cytoskeletal reorganization, stress responses and signaling suggesting that EVs containing these components could drive intercellular communication.

Third, TSG101 interacts with annexin A2 (ANXA2), a human protein implicated in small EV biogenesis. It, in turn, directly interacts with ANXA1 as well as three S100A Ca^2+^-binding proteins, GGCT and PHB implicated in cell proliferation. Most proteins in this subnetwork (83%) do not have yeast orthologs and none are implicated in protein degradation. Thus, we speculate that this may represent a synthetic EV protein sorting pathway in *S. cerevisiae* whereby yeast machinery (e.g. Vps23, the *S. cerevisiae* ortholog of human TSG101) may be capable of forming interactions with these cargoes. Together, these findings support the idea that EV cargo loading is not stochastic and may be guided by molecular connectivity to EV biogenesis pathways such as ESCRTs, and that *S. cerevisiae* provides a functionally conserved platform to uncover these mechanisms.

### Validation of results reveals candidate human scaffolds for engineering yeast EVs

To validate our findings, we selected seven candidate human proteins identified in *S. cerevisiae* EVs for follow-up analysis: Defensin A3 (DEFA3 or hBD-3), a small (5 kDa) antimicrobial peptide, was selected because it is relatively abundant in yeast EV samples based on label-free mass spectrometry analysis and is present in human EVs (Keshavarz Alikhani et al., 2021). We selected three human proteins with structural properties that may be valuable as a potential EV scaffolds facilitating surface display of engineered epitopes (Alvarez-Erviti et al., 2011; Nakase & Futaki, 2015): CLIC1 (Chloride Intracellular Channel 1), a 27-kDa protein that exists in soluble and membrane-associated states, exhibits glutaredoxin-like oxidoreductase activity in its soluble form, and can undergo redox-dependent structural transitions that promote membrane insertion and anion channel activity (Al Khamici et al., 2015; Littler et al., 2004; Stoychev et al., 2009) ; Orthologous to *S. cerevisiae* ERP5, human TMED9 (Transmembrane p24 Trafficking Protein 9) is a 27-kDa p24-family transmembrane protein involved in early secretory pathway trafficking and recently implicated in the capture and clearance of misfolded protein cargo during protein quality control (Ronzier & Satpute-Krishnan, 2025; Xiao et al., 2024); and QSOX1 (Quiescin Q6 Sulfhydryl Oxidase 1), an 82-kDa sulfhydryl oxidase that catalyzes disulfide bond formation during oxidative protein folding and can undergo proteolytic processing of its C-terminal transmembrane region to promote secretion of a soluble enzyme product (Chakravarthi et al., 2007; Rudolf et al., 2013).

We also selected three human proteins that potentially mediate interactions between ESCRTs and EV cargo proteins (based on Figure 5), are frequently identified in human EV samples (Vesiclepedia Top 100), and are not ubiquitin paralogs: Orthologous to Ssa2, an EV biomarker in *S. cerevisiae* (Logan et al., 2024), human HSPA8 (Heat Shock Protein A8, or HSC70) is a constitutively expressed 71-kDa molecular chaperone for protein folding, trafficking and degradation involved in chaperone-mediated autophagy (Kaushik & Cuervo, 2018); Orthologous to the *S. cerevisiae* EV protein Hsc82, human HSP90AB1 (Heat Shock Protein 90 Alpha B1) is a constitutively expressed 90-kDa molecular chaperone for protein folding and stabilization (Haase & Fitze, 2016); and human Annexin A2 (ANXA2; 36 kDa) is a calcium-dependent phospholipid binding protein thought to help mediate EV biogenesis (Gerke et al., 2024; Valapala & Vishwanatha, 2011; Varyukhina et al., 2022).

Each HsORF was individually cloned into the EVclo expression vector used for our screen, except GFP was replaced with mNeonGreen (mNG) to improve human fusion protein detection. We first confirmed fusion protein expression in *S. cerevisiae* transformants by flow cytometry and found that 41 to 95 % of cells contained mNG, depending on the HsORF expressed (Fig. 6a and Supplementary Fig. 4). We visualization these cells using confocal fluorescence microscopy (Fig. 6b) and found that all mNG-tagged proteins localized, at least in part, to intercellular puncta that may represent sites of ESCRT-mediated EV biogenesis. Membrane associated human proteins CLIC1, QSOX1 and ANXA2 seem to also be localized on plasma membranes. Western blot analysis confirmed that full-length fusion proteins were expressed, as prominent, anti-mNG reactive bands with predicted molecular weights were observed (Fig. 6c).

**Figure 6.**
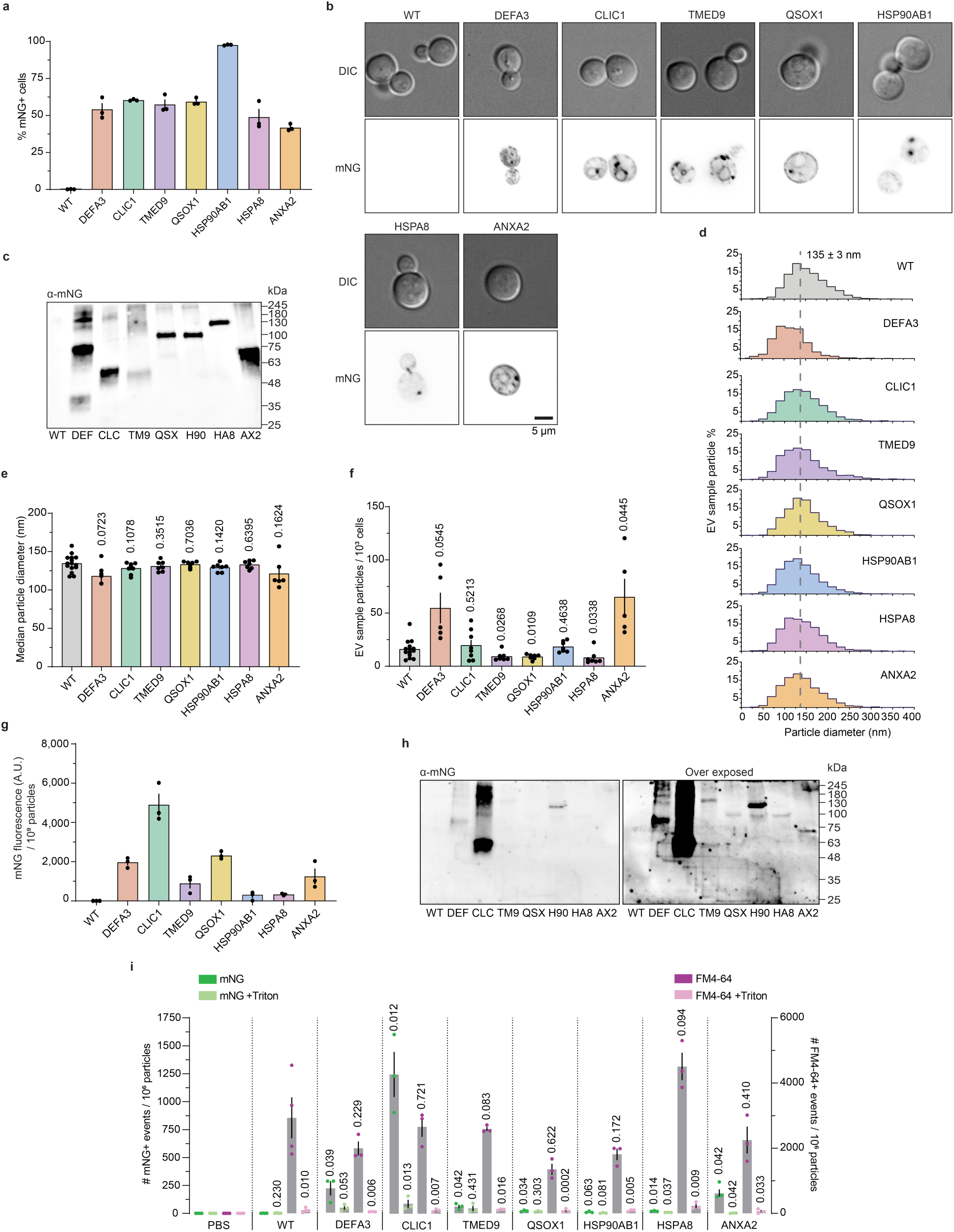
(previous page). Characterization of selected human EV cargo candidates expressed individually in *S. cerevisiae*. **a,** Percentage of mNeonGreen (mNG)-positive cells detected by flow cytometry. Candidate human EV proteins expressed individually in different strains and wild type (WT) are shown. Bars represent mean ± S.E.M. from n = 3 biological replicates. **b,** Representative DIC and inverted mNeonGreen (mNG) fluorescence micrographs of WT cells and cells expressing DEFA3, CLIC1, TMED9, QSOX1, HSP90AB1, HSPA8, or ANXA2 as C-terminal mNG fusions. Scale bar, 5 μm. **c,** Anti-mNG immunoblot of whole-cell lysates. Predicted molecular weights of mNG-fusion proteins are: DEFA3 (36.8 kDa; DEF), CLIC1 (53.5 kDa; CLC), TMED9 (53.9 kDa; TM9), QSOX1 (109.6 kDa; QSX), HSP90AB1 (110.0 kDa; H90), HSPA8 (97.5 kDa; HA8), or ANXA2 (65.2 kDa; AX2). **d – f,** Representative particle size distributions (**d**), mean particle diameter (**e**) mean particle numbers normalized to 1,000 cells (**f**), and mNG fluorescence normalized to 1 × 10^9^ particles (**g**) for EV samples isolated from WT and strains expressing human candidate EV proteins fused to mNG. Dashed line indicates mean particle diameter of WT (135 ± 3 nm). Bars represent mean ± S.E.M. from n = 3 biological replicates. **h,** Anti-mNG immunoblot of EV samples shown at two exposures. **i,** Nano-flow cytometry analysis of EV samples from each pool, normalized using NTA particle counts. Number of mNG-positive particles (green) and FM4-64-positive particles (magenta) are shown. Lighter shaded data points indicate samples treated with Triton X-100. PBS and WT served as a negative controls. Bars indicate means ± S.E.M.. n = 3 biological replicates. Statistical comparisons shown compare WT to each strain expressing a mNG-tagged candidate human protein with or without Triton X100 treament and were performed using a Welch’s *t*-test.

We next isolated EVs by ultrafiltration and size-exclusion chromatography from these strains and used nanoparticle tracking analysis to size and count particles in EV samples (Fig. 6d – f). Expressing these human cargo proteins had no effect on particle size (Fig. 6e; median 135 nm ± 3 nm diameter) suggesting that small EV or exosome biogenesis and release was unaffected. However, over-expression of mNG tagged DEF3A and ANXA2 seemed to increase EV titers, whereas TMED9, QSOX1 and HSPA8 seemed to decrease EV titers compared to EV yields from the parent yeast strain (Fig. 6f). To determine if nNG-tagged proteins were loaded into EVs, we measured EV sample fluorescence by fluorometry (Fig. 6g) and found that mNG-tagged CLIC1 showed the highest levels in EVs followed by DEF3A = QSOX1 > ANAX2 = TMED9 > HSP90AB1 = HSPA8. We conducted western blot analysis to confirm the presence of full length mNG-tagged fusion proteins in EV samples (Fig. 6h) and found that immunoreactivity was highest for CLIC1 followed by DEF3A = HSP90AB1 > ANAX2 = TMED9 > QSOX1 = HSPA8 with little free mNG detected suggesting that the fluorescence signal observed likely represents full-length human protein-mNG fusions.

To demonstrate that mNG-labeled proteins were found in lipid-bound particles representing EVs, we stained EVs in samples with the lipophilic dye FM4-64 and conducted nano-flow cytometry (Fig. 6i). We confirmed the presence of detergent (Triton X-100)-soluble, FM4-64 positive membrane-bound vesicles in all samples and detected the presence of detergent-soluble, mNG-positive particles isolated from strains expressing CLIC1 > DEFA3 = ANXA2 (Figure 6i). Samples prepared from strains expressing QSOX1, TMED9 and HSPA8 contained few detectable mNG positive particles, and mNG-HSP90AB1 was not detected in EV samples when analyzed using this method (Fig. 6i). To conclude, we are very confident that membrane associated human proteins CLIC1 and ANXA2 and the small human antimicrobial peptide DEFA3 are sorted into *S. cerevisiae* small EVs; human QSOX1, TMED9, HSP90AB1 and possibly HSPA8 may also be sorted into yeast EVs, but further analysis is needed to validate our preliminary findings.

## DISCUSSION

Herein, we establish *Saccharomyces cerevisiae* as a genetically tractable platform to study deeply conserved mechanisms responsible for protein cargo loading into small extracellular vesicles. Using a bespoke batch cloning protocol, we inserted 13,614 of 15,415 unique human ORFs into yeast expression plasmids, transformed them into *S. cerevisiae* and found that 3,849 were translated into proteins (Figures 1 – 3). Of these, 279 human proteins were detected within yeast EV samples; 72% of which have *S. cerevisiae* orthologs and most of these human proteins or their yeast orthologs were previously detected in EVs, including 31 of the top 100 proteins detected in human EVs (Figures 4 and 5). Analysis of interactions between these 279 human proteins and ESCRT components revealed three possible pathways for cargo loading into small EVs that involve ubiquitin, heat shock proteins or annexins (Figure 5). Validation studies confirm that at least three of seven human proteins tested – ANXA2, CLIC1 and DEFA3 – are sorted into *S. cerevisiae* small EVs (Figure 6), possibly representing scaffolds that can be used to engineer yeast EVs.

Mechanisms responsible for protein cargo sorting into small EVs are likely conserved from yeast to man. The selective recovery of 279 human proteins from a much larger pool of expressed human proteins argues against passive or stochastic incorporation and instead suggests that *S. cerevisiae* can recognize molecular features, interaction partners, or trafficking states that promote EV entry. This is particularly notable because yeast lacks certain mammalian-specific trafficking regulators, yet still incorporates many canonical human EV proteins, suggesting that modular cargo-recognition mechanisms are sufficiently conserved to operate in a heterologous host (Christ et al., 2017; van Niel et al., 2018). Orthology analysis further supports this interpretation, as most human proteins identified in yeast EVs have yeast counterparts, many of which are also detected in native yeast EV proteomes (Mencher et al., 2020; Zhao et al., 2019). Network analysis connected many of these proteins to ESCRT-associated pathways and revealed three possible cargo-sorting routes involving ubiquitin-linked proteins, heat shock chaperones, and annexin-associated proteins. These subnetworks may represent distinct but overlapping mechanisms by which human proteins are recruited into yeast EVs, either through direct ESCRT recognition, association with conserved protein complexes, or linkage to protein quality-control pathways. Although these interactions do not prove direct ESCRT-mediated sorting, they provide a testable framework for future studies. Targeted perturbation of ESCRT components, ubiquitin machinery, chaperones, and endosomal trafficking factors will be needed to determine which pathways are required for human protein incorporation into EVs and which operate independently of ESCRTs.

Fundamental EV functions may also be inferred from the conserved cargoes identified here. Among human EV proteins with yeast orthologs, 90 overlapped with proteins previously detected in native yeast EVs, and 20 were also among the most frequently reported human EV proteins in Vesiclepedia, including cytoskeletal regulators, glycolytic enzymes, redox proteins, chaperones, translation factors and 14-3-3 scaffolds (Logan et al., 2024; Pathan, et al., 2019). These shared cargoes point to conserved EV-associated functions in cytoskeletal remodeling, metabolism, proteostasis, stress adaptation and intracellular signaling (Kowal et al., 2016; Yáñez-Mó et al., 2015). Although some EV cargoes may act as individual functional proteins, many of the most conserved cargoes are components of larger molecular assemblies, suggesting that EVs may package functional modules rather than isolated proteins. Human proteins expressed in yeast may therefore associate with conserved yeast orthologous partners or enter EVs as part of mixed human–yeast complexes, as shown previously for humanized yeast protein assemblies such as the proteasome (Sultana et al., 2023). Conversely, several known human EV proteins lacking clear yeast orthologs were still recovered in yeast EVs, indicating that ancient vesicle-sorting pathways can accommodate later-evolving metazoan proteins. The validated incorporation of CLIC1, ANXA2 and DEFA3 further supports this idea, as these proteins represent membrane-associated, scaffold-like and antimicrobial cargo classes that could confer new properties on yeast EVs (Al Khamici et al., 2015; Panigrahi et al., 2022; Valapala & Vishwanatha, 2011). Thus, humanized yeast EVs may preserve ancient EV-associated functions while providing a route to introduce synthetic, human-specific activities into a simple eukaryotic vesicle system.

Finally, this work establishes *S. cerevisiae* as a versatile platform to dissect fundamental small EV biology and advance EV engineering. Its genetic tractability, rapid growth and compatibility with pooled screening make yeast well suited for testing how defined genes, pathways, protein domains and interaction motifs influence EV cargo selection. The human ORFeome screen therefore provides a resource for prioritizing candidate cargoes, scaffolds and sorting pathways that can be interrogated by targeted genetics, ESCRT perturbation, domain mapping and quantitative EV profiling. At the same time, the recovery and validation of human proteins in yeast EVs supports the use of *S. cerevisiae* as a chassis for engineering designer vesicles with defined molecular features. Proteins such as CLIC1, ANXA2 and DEFA3 provide starting points for developing cargo-loading, surface-display or functionalized EV strategies. Nonetheless, this study identifies candidate sorting relationships rather than causal mechanisms, and constitutive overexpression or mass spectrometry detection limits may influence the cargoes recovered. Future work combining pooled humanized yeast libraries with systematic gene deletion, cargo-domain mutagenesis, ortholog replacement and functional delivery assays should help define the rules that govern EV cargo selection. In this way, humanized yeast bridges basic EV biology and synthetic vesicle engineering, moving EV design from empirical optimization toward mechanism-guided construction.

## METHODS

### Yeast strains and reagents

All *Saccharomyces cerevisiae* strains used in this study are listed in Table 1. Unless otherwise noted, biochemical and growth reagents were purchased from Thermo Fisher Scientific (Burlington, Canada), BioShop Canada Inc. (Burlington, Canada), or Sigma-Aldrich (Oakville, Canada).

**Table 1.** *S. cerevisiae* strains used in this study.

| **Strains** | **Background genotype** | **Source** |
| --- | --- | --- |
| BY4741 | MATα *his3-Δ1 leu2-Δ0 met15-Δ0 ura3-Δ0* | (Huh et al., 2003) |
| hORFeome Pool 1-18-EGFP | BY4741, [pTDH3-HsORF-EGFP-tADH1 URA3] | This study |
| DEFA3-mNG | BY4741, [pTDH3-DEFA3-mNG-tADH1 URA3] | This study |
| CLIC1-mNG | BY4741, [pTDH3-CLIC1-mNG-tADH1 URA3] | This study |
| TMED9-mNG | BY4741, [pTDH3-TMED9-mNG-tADH1 URA3] | This study |
| QSOX1-mNG | BY4741, [pTDH3-QSOX1-mNG-tADH1 URA3] | This study |
| HSP90AB1-mNG | BY4741, [pTDH3-HSP90AB1-mNG-tADH1 URA3] | This study |
| HSPA8-mNG | BY4741, [pTDH3-HSPA8-mNG-tADH1 URA3] | This study |
| ANXA2-mNG | BY4741, [pTDH3-ANXA2-mNG-tADH1 URA3] | This study |

**Table 2.** Human proteins identified in EV samples and their *S. cerevisiae* orthologs.

| Human protein name | *S. cerevisiae* ortholog name(s) |
| --- | --- |
| ACTB | ACT1; ARP1; ARP6 |
| ACTN1 | SAC6 |
| AHCY | SAH1 |
| CFL1 | COF1 |
| EEF1A1 | GCD11; HBS1; SKI7; SUP35; TEF1; TEF2 |
| EEF2 | EFT1; EFT2; MEF1; MEF2; RIA1; SNU114 |
| EIF4A1 | FAL1; TIF1; TIF2 |
| ENO1 | ENO1; ENO2; ERR1; ERR2; ERR3 |
| GAPDH | TDH1; TDH2; TDH3 |
| HSP90AB1 | HSC82; HSP82 |
| HSPA1A | ECM10; KAR2; LHS1; SSA1; SSA2; SSA3; SSA4; SSB1; SSB2; SSC1; SSE1; SSE2; SSQ1; SSZ1 |
| HSPA8 | ECM10; KAR2; LHS1; SSA1; SSA2; SSA3; SSA4; SSB1; SSB2; SSC1; SSE1; SSE2; SSQ1; SSZ1 |
| PFN1 | PFY1 |
| PPIA | CPR1; CPR2; CPR3; CPR4; CPR5; CPR6; CPR7; CPR8 |
| PRDX1 | PRX1; TSA1; TSA2 |
| PRDX2 | PRX1; TSA1; TSA2 |
| RAN | GSP1; GSP2 |
| TPI1 | TPI1 |
| YWHAG | BMH1; BMH2 |
| YWHAQ | BMH1; BMH2 |

### Plasmids – CCSB Human ORFeome collection

The human ORFeome collection (HsORF) contains 15,465 human full-length cDNAs cloned into a Gateway recombinational entry vector (pDONR223). The native stop codon is removed to permit the expression of a tagged fusion protein. All 15,465 genes are segregated into 18 distinct pools of roughly 400 – 900 genes each. Clones were originally produced by the Center for Cancer Systems Biology (CCSB) at the Dana-Farber Institute (Rual et al., 2004).

### Plasmids – EVclo expression vector

HsORFs were ectopically expressed by introducing them into the EVclo destination vector (Bouffard et al., 2026), which includes the ampicillin resistance gene for antibiotic selection in *E. coli*, the 2µ yeast origin of replication for high copy number in *S. cerevisiae*, and the URA3 gene for auxotrophic selection in yeast. The EVclo destination vector has attR sites to enable Gateway cloning with the hORFeome entry clones, which contain attL sites. In the completed expression clone, the hORFeome gene is driven to high constitutive expression using the *S. cerevisiae* TDH3 promoter, and a 3’ EGFP (enhanced green fluorescent protein) or mNG (mNeon Green) gene in the destination vector produces a translational fusion protein with the human protein fused to a functional fluorescent protein for tracking.

### Humanized yeast strain creation by Gateway cloning

An en masse Gateway LR reaction was used to transfer the human ORFeome pool of entry clones (in pDONR223) into the destination vectors (EVclo): A 5 μl starter reaction containing 1 μl (150 ng) of human ORFeome pool in entry clone, 1 μl (150 ng) EVclo destination vector, 1 μl LR Clonase II Enzyme Mix, and 2 μl 1× TE buffer (Tris-EDTA) is incubated at room temperature overnight. A 5 μl reaction consisting of 1 μl (150 ng) destination vector, 1 μl LR Clonase II Enzyme Mix, 3 μl 1× TE buffer was then added to the starter reaction and incubated at room temperature overnight. This was repeated 3 additional times resulting in a 25 μl reaction ready for transformation after 5 days.

The entire reaction was added to 50 μl of DH5-alpha *E. coli* competent cells (New England Biolabs; Ipswich, USA; Cat #. C2987H) and transformed following the manufacturer’s instructions. Cells were plated on 245 x 245 mm square BioAssay dishes (Corning; Corning, USA) containing lysogeny broth (LB) with carbenicillin. Plates were then incubated at 37°C for 24 hours and transformants were scraped off for a plasmid extraction using a ZymoPURE II Plasmid Midiprep Kit (Zymo Research; Irvine, USA).

Competent *S. cerevisiae* cells (BY4741) were prepared using a Frozen-EZ yeast transformation II kit (Zymo Research; Irvine, USA) and stored at -80°C until use. 1 μg of extracted plasmid DNA is added to 1.8 x 10^6^ competent yeast cells for transformation following manufacturer instructions. Yeast were plated on 245 x 245 mm square BioAssay dishes (Corning; Corning, USA) containing synthetic complete medium lacking uracil (SC –URA). Plates were incubated at 30°C for 3 days. Transformants were scraped off and a pooled freezer stock was prepared (YPD medium with 25% glycerol).

### Plasmid sequencing

Pool 1 to 18 plasmid preparations are analyzed by Oxford Nanopore Technology sequencing (Plasmidsaurus; San Francisco, USA) until detection saturation was reached. Raw reads were analyzed with Geneious software (Dotmatics; Boston, USA).

### FACS enrichment of EGFP+ yeast cells

Glycerol stocks of pooled transformants or wild-type (negative control) yeast were used to inoculate 5 mL SC –URA or YPD seed cultures and incubated overnight at 30 °C. 500 µL of overnight seed culture was then added to 5 mL of fresh media and incubated for 4 hours at 30°C. After measuring culture density (OD_600nm_/mL), 5 OD units of seed culture were transferred to 1.5 mL microcentrifuge tubes, washed twice with 1 mL PBS (10,000 x g for 1 minute), and resuspended in 1 mL PBS (Cytiva; Marlborough, USA). 250 µL was then added to 750 µL PBS in a 5 mL FACS tube and EGFP-positive cells were collected using a Melody FACS instrument (BD Biosciences, Mississauga, Canada) with a rate of 1,000 – 2,000 events per second and cells were detected using a 488 nm laser and 527/32 nm bandpass filter. Freezer stocks (YPD with 25 % glycerol) were prepared using ∼1,300,000 events (yeast cells) collected from each sample.

### Flow cytometry of yeast cells

Glycerol stocks of EGFP+ pooled transformants or wild-type yeast used to inoculate 5 mL SC – URA or YPD seed cultures grown overnight at 30°C. 500 µL of seed culture was then added to 5 mL of fresh media and incubated for 4 hours at 30°C. After measuring culture density (OD_600nm_/mL), 5 OD units of cultures were transferred to 1.5 mL microcentrifuge tubes, washed twice with 1 mL PBS (10,000 x g for 1 min) and resuspended in 1 mL PBS. 250 µL of cell suspension was added to 750 µL of PBS in a 96 deep well microplate and then analyzed using Aurora flow cytometer and Spectroflo software (Cytek Biosciences; Freemont, USA) at the slow fluidics setting with 1 wash cycle between sampling each well, and particle (cells) were detected using a 488 nm laser and 525/40 nm bandpass filter. 20,000 events were measured for each sample.

### Live-cell fluorescence microscopy

Yeast colonies were used to inoculate a 5 mL SC -URA seed cultures and incubated overnight at 30°C. 500 µL of seed cultures were added to 5 mL of fresh media, incubated for 4 hours at 30°C, and culture density was measured (OD_600nm_/mL). 5 OD units of culture were transferred to 1.5 mL microcentrifuge tubes, washed twice with 1 mL PBS (10,000 x g for 1 minute), and resuspended in 100 µL PBS. 7 µL of cell suspension was transferred to a 15 mm x 25 mm microscope slide and covered with a 24 mm x 60 mm #1.5 glass coverslip. Samples were imaged using a Axio confocal microscope (Carl Zeiss; Toronto, Canada) outfitted with a CICERO (Crest Optics; Rome, Italy) spinning disk, 100x Plan Apo NA 1.46 oil immersion objective, LDI-5 laser launch (89 North; Williston, USA) and Flash 4.0 LT Plus sCMOS camera (Hamamatsu Photonics; Bridgewater, USA; model #C11440-42U30) with 16-bit depth, 2048 x 2048 pixels. Brightfield images were acquired using white transmitted light with a 100 ms exposure time. A 488 nm laser (60 % power) and 525/50 nm emission filter were to acquire EGFP fluorescent images at 100 ms exposure. Volocity software (vs 7.0.0; Quorum Technologies Inc.; Puslinch, Canada) was used for image acquisition.

### Extracellular vesicle isolation

Seed cultures were prepared by inoculating from a glycerol freezer stock of yeast into 15 mL of YPD medium followed by incubation at 30°C for 8 hours in an orbital shaking incubator at 200 rpm. Seed cultures were used to inoculate 1 L of YPD medium in 2 L flasks incubated overnight at 30°C in an orbital shaking incubator at 200 rpm. When cultures reached densities of 8 – 12 OD_600nm_/mL, they were transferred to two 500 mL PPCO centrifuge bottles (ThermoFisher Scientific; Cat# 21020–050) and harvested by centrifugation (3000 × *g*, 5 minutes) at room temperature. Supernatants were discarded, and cell pellets were recovered and washed twice with PBS. Washed cell pellets were then exposed to heat conditioning at 42°C for 15 minutes in a water bath, immediately resuspended in 25 mL 0.1μm filtered PBS, and incubated for an additional 30 min at 42°C. Concentrated yeast samples in PBS were transferred to 50 mL polycarbonate centrifuge bottles (Beckman Coulter; Mississauga, Canada; Cat# 357002) pre-chilled on ice. All subsequent steps were carried out on ice or at 4°C.

Yeast samples were subjected to centrifugation (15,000 × *g*, 15 minutes), supernatants (containing EVs) were recovered, filtered using pre-chilled 0.22 μm filters to remove cell debris, and transferred to a 15 ml 100 kDa Amicon centrifugal filtration unit. Samples were then subjected to centrifugation (3,200 × *g* at 4°C) until volumes were ∼150 μL, and then transferred to sterile 1.5 mL microcentrifuge tubes on ice prior to isolation via size exclusion chromatography (SEC). SEC was performed using 150 μl qEV columns (IZON Science LTD; Medford, USA) and fractions 6 to 9 were pooled for downstream EV analysis.

### EGFP Immunoprecipitation (EGFP-IP)

To isolate EGFP–fusion proteins from whole cell lysates, 100 OD_600nm_ units of yeast transformants grown in liquid SC medium to mid-log phase were collected by centrifugation (3,500 × g, 5 minutes, room temperature). All subsequent steps were performed at 4°C or on ice. Cell pellets were resuspended in IP lysis buffer (50 nM HEPES-KOH, 150 mM KOAc, 2 mM MgOAc, 1 mM CaCl_2_) supplemented with protease inhibitor cocktail (Abcam; Waltham, USA) and 0.1% digitonin. Acid-washed glass beads were added and yeast were lysed using a cell disruptor (Scientific Industries; Bohemia, USA). Equal parts of IP lysis buffer supplemented with protease inhibitor cocktail and 1.9 % digitonin were then added, followed by incubation at 4°C on a nutator for 45 minutes, and centrifugation (13,000 x g, 5 minutes) to collect the supernatant representing the whole cell lysate. 30 μL GFP-Trap beads (Chromotek GFP-Trap Agarose; Proteintech; Rosemont, USA) were washed with 0.1% digitonin IP lysis buffer, added to whole cell lysates, and incubated for 1 hour on a nutator. Samples were washed 4 times with 0.1% digitonin in IP lysis buffer, prior to addition of 30 μl elution buffer (150 mM Tris pH 6.8, 6 M Urea, 6% SDS, 40% Glycerol, 100 mM DTT, 0.01% Bromophenol blue), incubation at 95°C for 5 minutes and collection of eluates.

To isolate EGFP-fusion proteins from EV samples, EV samples prepared by SEC added to 2.5% digitonin IP lysis buffer and incubated on a nutator at 4°C for 45 minutes. 30 μL of washed EGFP-Trap beads were then added and samples were incubated for 1 hour on a nutator, and washed 4 times with 0.1% digitonin IP lysis buffer. Beads were then resuspended in 30 μl elution buffer, incubated at 95°C for 5 minutes, and eluates were collected.

### Liquid chromatography-tandem mass spectrometry (LC-MS/MS)

For proteomic analysis by LS-MS/MS, immunopurified EGFP-tagged human proteins from cell lysates or EV samples were loaded on a TGX gel (Bio-Rad Mini-protean 10% Tris-glycine gel) and the expected mass range of each pool was excised. Gel fragments were added to 50 mM ammonium bicarbonate with 10 mM TCEP (Tris(2-carboxyethyl)phosphine hydrochloride), and vortexed for 1 hour at 37°C. Chloroacetamide was added for alkylation to a final concentration of 55 mM. Samples were vortexed for another hour at 37°C. 1 µg trypsin was added, and digestion was performed for 8 hours at 37°C. Samples were dried and solubilized in 4% formic acid (FA). Peptides were loaded and separated on a reversed-phase column (150 μm i.d. by 200 mm) with a 56-min gradient from 10 to 30% ACN-0.2% FA and a 600 nL/min flow rate on an Easy nLC-1000 connected to an Orbitrap Fusion (Thermo Fisher Scientific; San Jose, USA). Each full MS spectrum acquired at a resolution of 120,000 was followed by tandem-MS (MS-MS) spectra acquisition on the most abundant multiply charged precursor ions for a maximum of 3s. Tandem-MS experiments were performed using collision-induced dissociation (CID) at a collision energy of 30%. Data were processed using PEAKS X Pro software (Bioinformatics Solutions; Waterloo, ON) using the UniProt human protein database (20,366 entries; The UniProt Concostrium, 2025). Mass tolerances on precursor and fragment ions were 10 ppm and 0.3 Da, respectively. The fixed modification was carbamidomethyl (C). Variable selected posttranslational modifications were oxidation (M), deamidation (NQ), phosphorylation (STY) along acetylation (N-ter). The data was visualized with Scaffold 5.0 software (Proteome Software, Inc.; Portland, USA) with a protein threshold of 99%, at least 2 peptides identified and a false-discovery rate of 1% for peptides. Proteomics data was viewed with Scaffold 5.0 and further analyzed with FUNRICH software (Pathan et al., 2015).

### Transmission electron microscopy

EV samples were fixed by diluting them 1:1 with 2.5% glutaraldehyde in 0.1 M sodium cacodylate. Fixed EVs (5 μL) were then drop cast onto glow-discharged carbon-coated grids and allowed to adsorb for 5 minutes at room temperature. EV-bound grids were washed twice with glycine and then rinsed four times with ultrapure pure water. Mounted EVs were then negatively stained with 1% phosphotungstic acid (1 minute, room temperature), blotted with filter paper, and airdried (1 hour, room temperature). Grids were then imaged at 80−120 kV using a Talos L120C transmission electron microscope (Thermo Fisher Scientific; Toronto, Canada).

### Particle (EV) size and concentration measurements

To measure particle (EV) size and concentrations, we conducted nanoparticle tracking analysis (NTA) using a ZetaView nanoparticle tracking analysis instrument (Particle Metrix; Ammersee, Germany) with software version 8.0.5.14 SP7. EV samples were diluted to achieve 50 to 200 particles per frame in sterile, tissue culture-grade PBS. 1 mL of diluted EV samples was manually loaded with a 1 mL syringe and samples were slowly injected. ZetaView instrument settings were as follows: Temperature (25°C), laser λ (488 nm), filter λ (scatter) sensitivity (80), shutter (100), frame rate (30), cycles (3), positions (11), and trace length (30).

### Western blot analysis

Whole cell lysates were prepared by collecting 1 OD_600nm_ unit of yeast cells grown in liquid SC medium to mid-log phase using centrifugation (3,500 × g, 5 minutes, room temperature). Pellets were resuspended in boiling buffer (1.5 M Tris pH 8.5, 0.5 M EDTA, 10% SDS) supplemented with protease inhibitor cocktail ((Abcam; Waltham, USA) and acid-washed glass beads were added. Samples were then subjected to 5 minutes in a cell disruptor (Scientific Industries; Bohemia, USA). Sample loading buffer (150 mM Tris pH 6.8, 6 M Urea, 6% SDS, 40% Glycerol, 100 mM DTT, 0.01% Bromophenol blue) was added, samples were disrupted again and incubated at 95°C for 5 min prior to analysis by SDS-PAGE.

EV samples collected after SEC were concentrated to roughly 15 µL using a 0.5 mL 10 kDa MWCO Amicon filter (Millipore; Oakville, Canada; Cat# 501024). 5 µl of 4x Laemmli sample buffer (Bio-Rad; Mississauga, Canada; Cat#1610747,) supplemented with protease inhibitor cocktail (Abcam; Waltham, USA; Cat# ab271306) and 50 mM DTT was added, and the sample was incubated at 95°C for 5 minutes prior to loading onto a gel.

Protein samples were separated by Mini-PROTEAN 10 % Tris-glycine protein gel (Bio-Rad; Mississauga, Canada), transferred to PVDF membranes using a Trans-Blot Turbo transfer system (Bio-Rad; Mississauga, Canada), and incubated with 5 % skim milk in 1x TBST on a nutator for 1 hour at room temperature. Membranes were then transferred to 5 % milk in 1x TBST containing anti-GFP mouse monoclonal antibody (MilliporeSigma; Mannheim, Germany; Cat#11814460001) at 1:1,000 dilution and incubated on a nutator at 4°C for 24 hours. Membranes were washed 5 times with 1x TBST, incubated for 45 minutes on a nutator at room temperature with 5 % milk in 1x TBST containing horseradish peroxidase-labeled affinity purified goat anti-mouse IgG antibody (SeraCare; Milford, USA; Cat#5450-0011) at 1:10,000 dilution, and then washed an additional 5 times with 1x TBST. Chemiluminescence of stained membranes was detected using a GE Amersham Imager 600 instrument (GE HealthCare; Piscataway, USA) and Amersham ECL Select Detection Reagent.

### Nano-flow cytometry of EV samples

Isolated EVs (5 x 10^9^) were stained with FM 4-64 ((*N*-(3-Triethylammoniumpropyl)-4-(6-(4-(Diethylamino) Phenyl) Hexatrienyl) Pyridinium Dibromide); Thermo Fisher Scientific; Cat#T13320) to a final concentration of 20 μM and incubated for 15 minutes at 30°C. Removing unbound dye was completed using the 70 nm qEV single SEC column (IZON Science LTD; Medford, USA) according to the manufacturer’s instructions. Stained EV fractions were kept at 4°C no longer than a day before nanoflow cytometry analysis.

EV sample concentrations were determined using NTA as described above, and samples were diluted to 3–5 × 10⁷ particles/mL in 500 µL PBS prior to flow cytometry. EVs from pooled transformants expressing EGFP were analyzed on a CytoFLEX nano-flow cytometer (Beckman Coulter; Mississauga, Canada) at a flow rate of 10 µL/min for 10 min, with water washes between samples. GFP fluorescence was excited using the 488 nm blue laser with emission collected in the FITC channel (525/40 nm bandpass), while FM4-64 fluorescence was excited using the 561 nm yellow-green laser with emission collected in the PC7 channel (780/60 nm bandpass). EV from clones expressing mNeonGreen were analyzed on an Aurora Northern Lights spectral flow cytometer (Cytek Biosciences; Freemont, USA) at 10 µL/min for 2 minutes per sample, with an abort rate maintained below 5% and water washes between samples. mNeonGreen fluorescence was excited using the 488 nm blue laser with emission collected in the B2 channel (525/40 nm bandpass), while FM4-64 fluorescence was excited using the 561 nm yellow-green laser with emission collected in the B9 channel (697/20 nm bandpass).

Detergent sensitivity controls were performed by adding Triton X-100 to EV samples at a final concentration of 1%, followed by vortexing and incubation at room temperature for at least 30 min before re-acquisition using identical settings. Gates for EGFP⁺/mNeonGreen⁺ and FM4-64⁺ events were established using wild-type EV controls, and fluorescence-positive event counts were normalized to the number of particles assayed based on NTA-derived particle concentrations for the acquisition volume.

### Bioinformatics and data analysis

Orthologous relationships between human and *Saccharomyces cerevisiae* were identified using the DIOPT (DRSC Integrative Ortholog Prediction Tool; Hu et al., 2011). Gene Ontology (GO) term annotations were obtained from gProfiler (Kolberg et al., 2020). Functional enrichment analyses were conducted using custom R scripts (version 2024.12.0+467), and network diagrams were visualized using Cytoscape (Shannon et al., 2003) with the STRING extension for protein-protein interaction networks. For proteomic analysis all human keratin isoforms were excluded, as they are assumed to be potential contaminants. For micrograph processing, all images were rotated, cropped, adjusted for brightness and contrast, inverted, annotated, and/or had color channels merged using Fiji/ImageJ (version 1.54f; Schindelin et al., 2012) and Adobe Illustrator and Photoshop CC software (Adobe Inc.; San Jose, USA). Data compilation and statistical analysis were performed using Microsoft Excel. All figures were prepared using Adobe Illustrator CC, and statistical significance of graphs was evaluated using GraphPad Prism (version 10.4.1; Dotmatics; Boston, USA), with specific tests and statistical parameters described in the figure legends.

## DATA AVILABILITY STATEMENT

Data that support findings of this study are available from the corresponding author upon reasonable request.

## ACKNOWLWDGEMENTS

We thank Dr. Nadim Tawil, Laura Montermini, and Hélène Pagé-Veillette at the Centre for Applied Nanomedicine at the McGill University Health Centre for assistance with nanoflow cytometry. We thank David Liu and Kelly Sears at the Facility for Electron Microscopy Research at McGill University for their assistance with transmission electron microscopy imaging.

Proteomics analysis was conducted with assistance from Dr. Éric Bonneil at the Institute for Research in Immunology and Cancer (IRIC), Université de Montréal. Dr. Christopher Law assisted with fluorescence microscopy conducted in the Centre for Microscopy and Cell Imaging at Concordia University. J.T. received a fellowship from the Synthetic Biology Applications training program funded by the Natural Sciences and Engineering Research Council of Canada (NSERC). This research was supported by research grant 2022-PR-298412 to C.L.B. and A.H.K from the Fonds de recherche du Québec – Nature et technologies

## AUTHOR CONTRIBUTIONS

J.T. – Conceptualization, Formal Analysis, Investigation, Methodology, Validation, Visualization, Writing; S.D. – Formal Analysis, Investigation, Visualization; A.H.K. – Conceptualization, Funding Acquisition; C.L.B. – Conceptualization, Funding Acquisition, Methodology, Project Administration, Resources, Supervision, Visualization, Writing.

## MATERIALS & CORRESPONDENCE

Correspondence and material requests should be sent to Christopher L. Brett.

For social media (265 characters):

By expressing the human ORFeome in *S. cerevisiae*, Trani et al. observe conserved mechanisms underlying extracellular vesicle (EV) cargo sorting. This study establishes humanized yeast to dissect EV biology and possibly engineer EVs for broad applications.

