## Supplementary Materials for "Human ORFeome expression in *S. cerevisiae* to better understand extracellular vesicle biology"

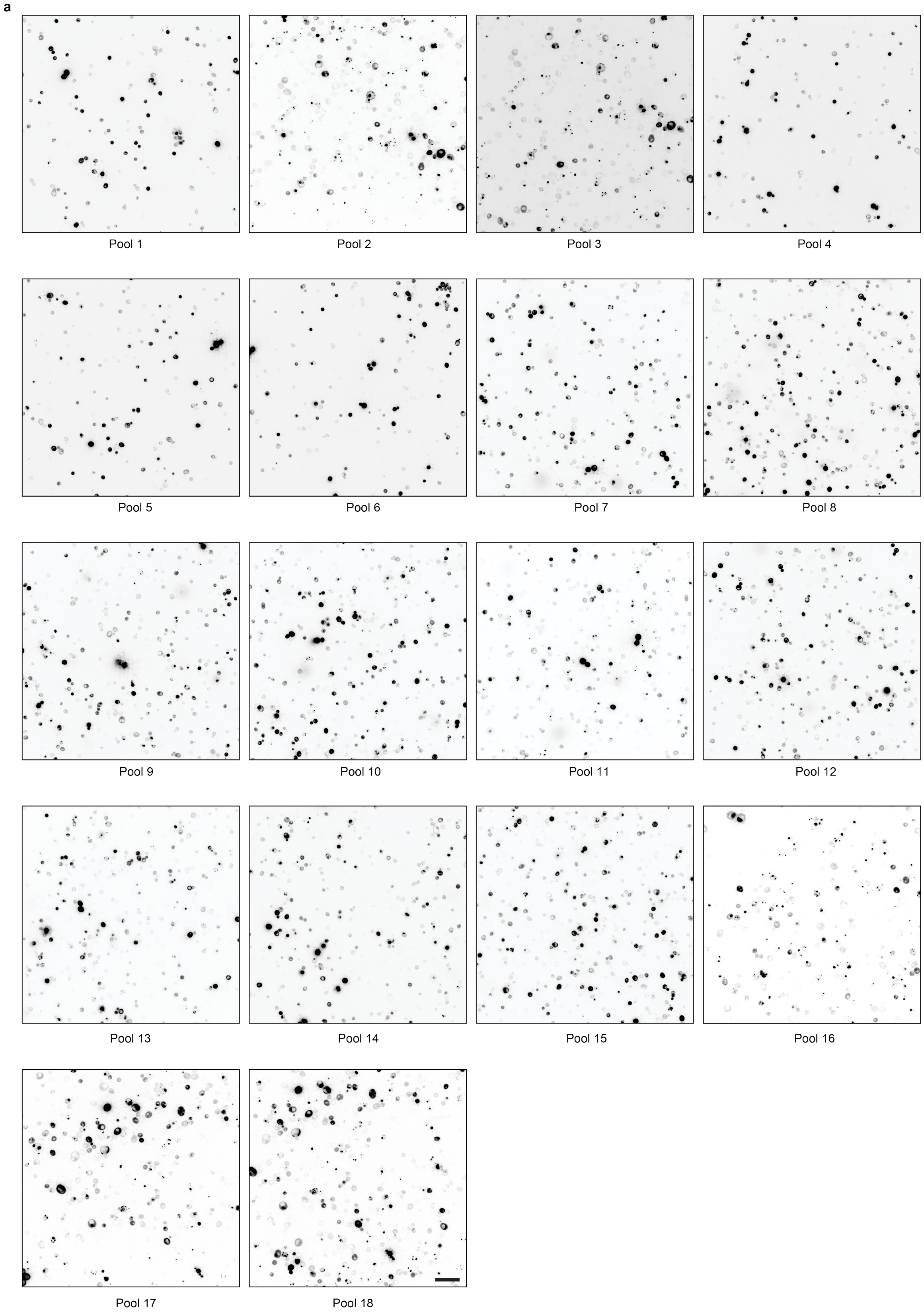

**Supplementary Figure 1 (previous page). Representative inverted GFP fluorescence micrographs of pooled humanized *S. cerevisiae*.**

**a,** Representative inverted GFP fluorescence micrographs of *S. cerevisiae* transformants from pools 1–18. n = 3 biological replicates. Scale bar, 20 µm.

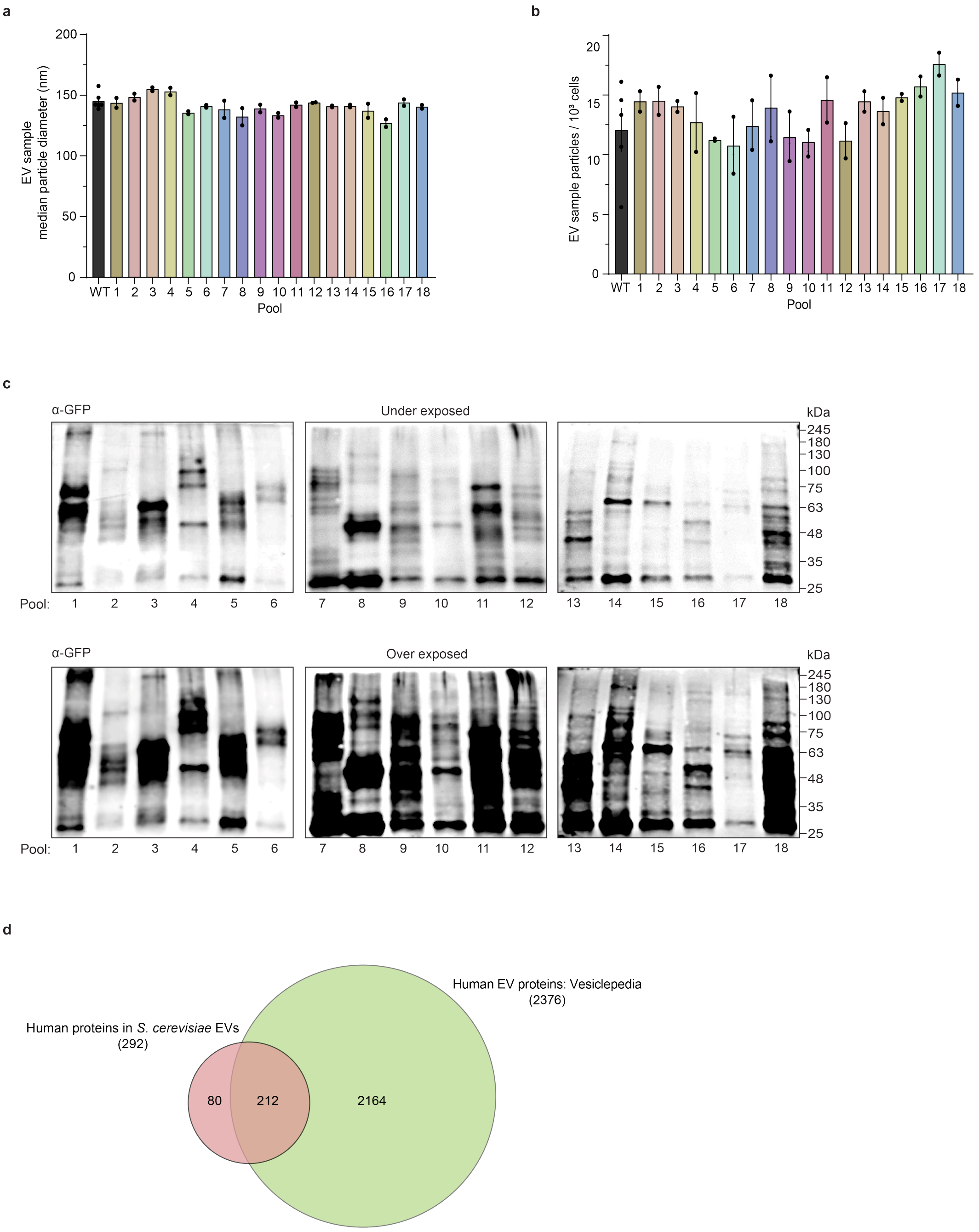


**Supplementary Figure 2. Characterization of extracellular vesicles isolated from pooled humanized *S. cerevisiae*.**

**a,** Median particle diameter of EV samples isolated from wild-type (WT) *S. cerevisiae* or yeast strains expressing human ORFeome pools 1–18 measured by nanoparticle tracking analysis (NTA). **b,** EV particle abundance normalized to 1,000 cells for WT and pools 1–18. **c,** Representative anti-GFP immunoblots of EV samples collected from pools 1–18, shown at two exposures to visualize high- and low-abundance GFP-tagged human proteins. Molecular weight markers are indicated in kDa. n = 3 biological replicates. **d,** Venn diagram showing overlap between human proteins identified in *S. cerevisiae* EVs and human EV proteins catalogued in Vesiclepedia. Of the 292 human proteins detected in yeast EV samples, 212 overlapped with Vesiclepedia-listed human EV proteins. Bars represent mean ± S.E.M. from *n* = 2 biological replicates.

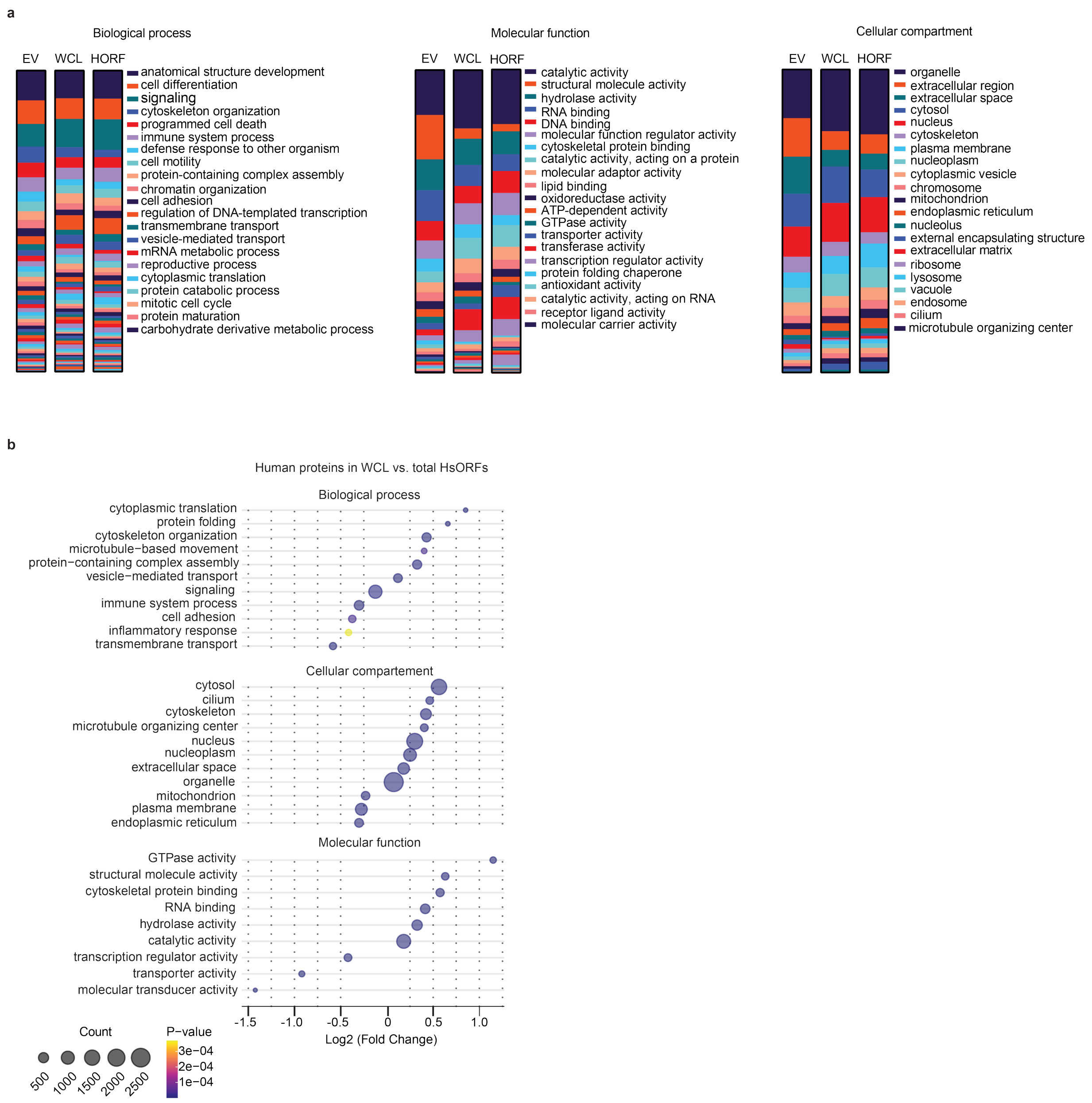


**Supplementary Figure 3. Gene Ontology annotation and enrichment analysis of human proteins identified in pooled humanized *S. cerevisiae***

**a,** Gene Ontology (GO) term distributions for human proteins found in EV samples, whole cell lysates (WCLs), or the HsORFeome library (HORF), shown for Biological Process, Molecular Function, and Cellular Compartment categories. **b,** GO enrichment and depletion analysis of human proteins identified in WCLs relative to the total HsORFeome input library, shown for Biological Process, Molecular Function, and Cellular Compartment categories. Dot size indicates protein count and color indicates $P$ value. Only categories with significant enrichment or depletion are shown.

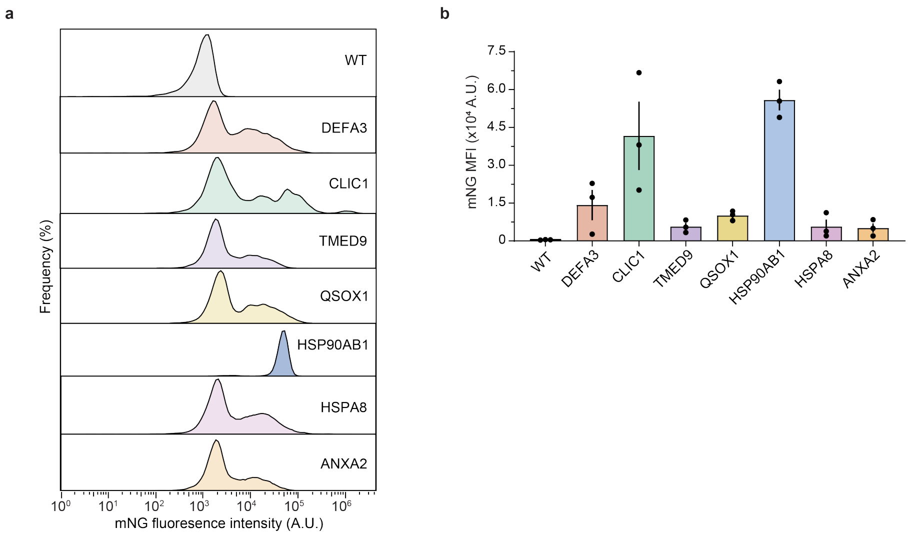


**Supplementary Figure 4. Flow cytometry analysis of mNeonGreen-tagged human protein expression in *S. cerevisiae*.**

**a**, Flow cytometry histograms showing mNeonGreen (mNG) fluorescence distributions for WT cells and cells expressing human DEFA3, CLIC1, TMED9, QSOX1, HSP90AB1, HSPA8, or ANXA2 as C-terminal mNG fusions. **b**, mNG median fluorescence intensity (MFI) measured by flow cytometry. Bars represent mean ± S.E.M. from n = 3 biological replicates.

DESCRIPTION OF ADDITIONAL SUPPLEMENTARY FILES

**Supplementary Information 1**

Sequencing data used to identify HsORFs cloned into pooled expression plasmids and to prepare Fig. 1b–e.

**Supplementary Information 2**

Human proteins identified by mass spectrometry in whole cell lysates (WCLs; Fig. 2h) or extracellular vesicle (EV) samples (Fig. 3e) from pools of yeast transformants.

**Supplementary Information 3**

Yeast orthologs of human proteins identified in EV samples compared to previously reported yeast EV proteins (Logan et al., 2024) and Vesiclepedia Top 100. Data was used to prepare Figs. 4 and 5.

**Supplementary Information 4**

Dataset supporting the human protein-protein interaction network shown in Fig. 5.
